# Advancing admixture graph estimation via maximum likelihood network orientation

**DOI:** 10.1101/2021.02.02.429467

**Authors:** Erin K. Molloy, Arun Durvasula, Sriram Sankararaman

## Abstract

**Motivation:** Admixture, the interbreeding between previously distinct populations, is a pervasive force in evolution. The evolutionary history of populations in the presence of admixture can be modeled by augmenting phylogenetic trees with additional nodes that represent admixture events. While enabling a more faithful representation of evolutionary history, *admixture graphs* present formidable inferential challenges, and there is an increasing need for methods that are accurate, fully automated, and computationally efficient. One key challenge arises from the size of the space of admixture graphs. Given that exhaustively evaluating all admixture graphs can be prohibitively expensive, heuristics have been developed to enable efficient search over this space. One heuristic, implemented in the popular method TreeMix, consists of adding edges to a starting tree while optimizing a suitable objective function.

**Results:** Here, we present a demographic model (with one admixed population incident to a leaf) where TreeMix and any other starting-tree-based maximum likelihood heuristic using its likelihood function is *guaranteed* to get stuck in a local optimum and return an incorrect network topology. To address this issue, we propose a new search strategy that we term maximum likelihood network orientation (MLNO). We augment TreeMix with an exhaustive search for a MLNO, referring to this approach as OrientAGraph. In evaluations including previously published admixture graphs, OrientAGraph outperformed TreeMix on 4/8 models (there are no differences in the other cases). Overall, OrientAGraph found graphs with higher likelihood scores and topological accuracy while remaining computationally efficient. Lastly, our study reveals several directions for improving maximum likelihood admixture graph estimation.

**Availability:** OrientAGraph is available under the GNU General Public License v3.0 on Github (https://github.com/sriramlab/OrientAGraph).

## 1 Introduction

Admixture, the exchange of genes between distinct populations, has emerged as an important process in shaping genetic diversity within and between populations. Evidence of gene flow events in many species, including humans, (Green *et al*., 2010), wolves (Pilot *et al*., 2019) and butterflies (Edelman *et al*., 2019), has led to increasing interest in estimation of the demographic history of populations while accounting for admixture.

The demographic history of populations in the presence of admixture can be naturally modeled by augmenting phylogenetic trees with additional nodes that represent admixture events. These *admixture graphs* cannot be directly observed and need to be estimated from genomic data. Admixture graphs model changes in allele frequencies as a Wright-Fisher process, parametrized by the pair (*N*, **Θ**), where *N* is a *phylogenetic network* on a set *S* of populations (Definition 1) and **Θ** is a real-valued vector describing parameters associated with the edges of *N*, namely branch lengths in units of genetic drift and admixture proportions. The objective of admixture graph inference is to estimate (*N*, **Θ**) from genetic variation data measured across the populations in *S*. Many popular methods utilize summary statistics, such as the vector ***X*** of *f*-statistics, computed from allele frequencies in each of the *S* populations.

One of the most widely used tools, `qpGraph` (Patterson *et al*., 2012), takes ***X*** as well as a network topology *N* as input and then fits the numerical parameters **Θ**. When the number of populations is very small, it is possible to evaluate the likelihood of all network topologies with a fixed number of admixture events (e.g. using the `admixturegraph` R package from Leppälä*et al*., 2017). Otherwise, the researcher must propose potential networks and evaluate them individually (see Harney *et al*., 2021 for discussion).

Brute force and manual procedures quickly become intractable, as the number of possible network topologies grows super-exponentially in the number of populations (McDiarmid *et al*., 2015). This has motivated the development of heuristics for estimating admixture graphs from *f*-statistics. For example, `MixMapper` (Lipson *et al*., 2013, 2014) is a pipeline in which the user identifies a set of unadmixed populations *R ⊆ S*, estimates a submodel (i.e. a population tree) on *R*, and then sequentially adds populations in *S \ R* to the submodel. This approach works best when there are only a few admixed populations and can be viewed as a “semi-automated” version of `qpGraph` (Lipson, 2020).

A recent approach to automating admixture graph inference, called `miqograph` (Yan *et al*., 2020), is unique from prior methods in that it simultaneously estimates the network topology *N* and numerical parameters **Θ**, requiring the user to specify some additional parameters. `miqograph` is similar to `admixturegraph` in that it is guaranteed to find the highest scoring admixture graph with a user-specified number of admixture events; however, unlike `admixturegraph`, the network space evaluated by `miqograph` is constrained to networks in which all admixture nodes are either a leaf node or are the parent of an admixture node (see Yan *et al*., 2020 for further discussion).

To the best of our knowledge, the only method that is fully automated and (at least in principle) allows for the greatest flexibility in exploring network topologies is `TreeMix` (Pickrell and Pritchard, 2012). The only topological constraint imposed by `TreeMix` is that *N* is *tree-based*, meaning that it can be drawn as a tree annotated with additional edges representing gene flow (Francis and Steel, 2015; see Definition 4). At a high level, `TreeMix` uses a maximum likelihood (ML) search heuristic that operates by searching for an ML population tree *T* on *S* and then adding edges sequentially provided the edge additions improve a measure of model fit (hill climbing). We refer to methods that operate in this fashion as starting-tree-based maximum likelihood (STB-ML) methods. Empirical evidence suggests that `TreeMix` can be biased by its starting tree, especially when many populations are admixed (Lipson *et al*., 2013), so researchers may be wary of relying solely on `TreeMix`, despite its scalability relative to other approaches (Lipson, 2020).

In this paper, we consider the problem of admixture graph estimation from *f*-statistics using STB-ML methods. To explore the limitations of this class of methods, we study a simple admixture graph (*N*^*\**^, **Θ**^*\**^) with just one admixture event incident to a leaf. Given the *f*-statistics implied by the true model (*N* ^*\**^, **Θ**^*\**^) (which is equivalent to assuming that the number of independent sites goes to infinity), any STB-ML method that succeeds in finding the ML tree (which in this case is also the Neighbor Joining tree) as its starting tree, is guaranteed to get stuck in a local optimum and return an incorrect network topology *N*′. Our case study highlights that STB-ML methods can be inconsistent even under a demographic model with a single admixture event.

Examining our case study in more detail, we observe that the incorrect topology *N*′ can be transformed into the correct topology *N*^*\**^ simply by changing which population is admixed and redirecting the edges of *N*′ accordingly. This graph transformation is performed by researchers manually searching for the admixture graph topology using `qpGraph` (Lipson, 2020) and is also related to recent theoretical results on phylogenetic network orientation by Huber *et al*. (2019). Utilizing the terminology of Huber *et al*. (2019), we say that *N*′ and *N*^*\**^ are two different *orientations* of the same undirected network (Definition 5), with *N*^*\**^ yielding in a higher likelihood score than *N*′. While Huber *et al*. (2019) look at network orientation from a graph theoretic perspective (e.g. addressing questions such as: is the orientation of an undirected network uniquely determined by the position of the root and admixed populations?), here we explore network orientation as a search strategy, leading us to propose the *maximum likelihood network orientation* (MLNO) problem.

To evaluate the utility of MLNO for improving the accuracy of STB-ML methods based on *f*-statistics, we augment `TreeMix` with an exhaustive search for a MLNO after every edge addition. In line with our theoretical expectations, this approach, which we refer to as `OrientAGraph`, recovers the correct network in our motivating case study, whereas the original `TreeMix` method does not. We also benchmark `OrientAGraph` against `TreeMix` and `miqograph` on 7 model admixture graphs, 6 of which were estimated from biological data sets in recent studies. We find that `OrientAGraph` improves the accuracy of the original `TreeMix` method and, in two scenarios, is more accurate than `miqograph`. The first scenario occurs when an admixed population is not incident to a leaf (as this violates the topological constraints employed by `miqograph`); the second scenario occurs when the data set is large enough (*e*.*g*., 10 populations and 2 admixture events) so that `miqograph` is unable to solve its problem to optimality within our specified maximum running time. We conclude by discussing future directions as well as the potential utility of MLNO to STB-ML methods that take estimated gene genealogies as input (e.g. Yu *et al*., 2014; Wen *et al*., 2018; Wu, 2020).

## 2 Terminology and Background

Throughout this paper, we discuss results for phylogenetic networks and the approximate likelihood function computed by `TreeMix` (Pickrell and Pritchard, 2012); the relevant background is provided in this section.

### 2.1 Phylogenetic Networks

We use the term *phylogenetic network* to refer to a graphical object, which we now define.

#### Definition 1

(Phylogenetic Network). A *phylogenetic network N* is a triplet (*n, S, ϕ*), where *n* is a graph, *S* is a set of labels (typically denoting species or populations), and *ϕ* is a bijection mapping the leaves (i.e. vertices with out-degree 0) of *n* to the labels in *S. N* is *directed*, meaning that *n* is a directed acyclic graph with a directed path between the root (i.e. a special vertex with in-degree 0) and all other vertices in *n* and without any parallel arcs or self-loops. For convenience, we typically do not make an explicit distinction between a phylogenetic network *N* and its graph *n*. Instead, we say that *N* is a network on *S* and denote its vertices as *V* (*N*), edges as *E*(*N*), and leaves as *L*(*N*).

Henceforth, we assume that *N* is *binary*, meaning the root has out-degree 2, leaf vertices have indegree 1, and all other vertices, referred to as internal vertices, have in-degree and out-degree summing to 3. This is consistent with Francis and Steel (2015), Huber *et al*. (2019), and much of the literature on phylogenetic networks. However, we use the terms admixture or gene flow instead of reticulation or hybridization. Specifically, we refer to any internal vertex with in-degree *≥* 1 as an *admixture node*. An *admixture edge* is an arc whose head (target vertex) is an admixture node; all other arcs are *tree edges*. A directed phylogenetic network with zero admixture nodes is a directed *phylogenetic tree*.

`TreeMix` and related methods search the space of directed phylogenetic networks using operations, such as *edge additions* and *tail moves* (Figure 2). Such operations are *legal* if they produce a directed phylogenetic network. We refer to the set of networks that can be created by performing one legal edge addition to *N* as the edge addition *neighborhood* of *N*; we use similar terminology for other operations, such as tail moves. It is worth noting that the space of all directed phylogenetic networks is connected under tail moves (Janssen *et al*., 2018; Gambette *et al*., 2017).

**Figure 1:**
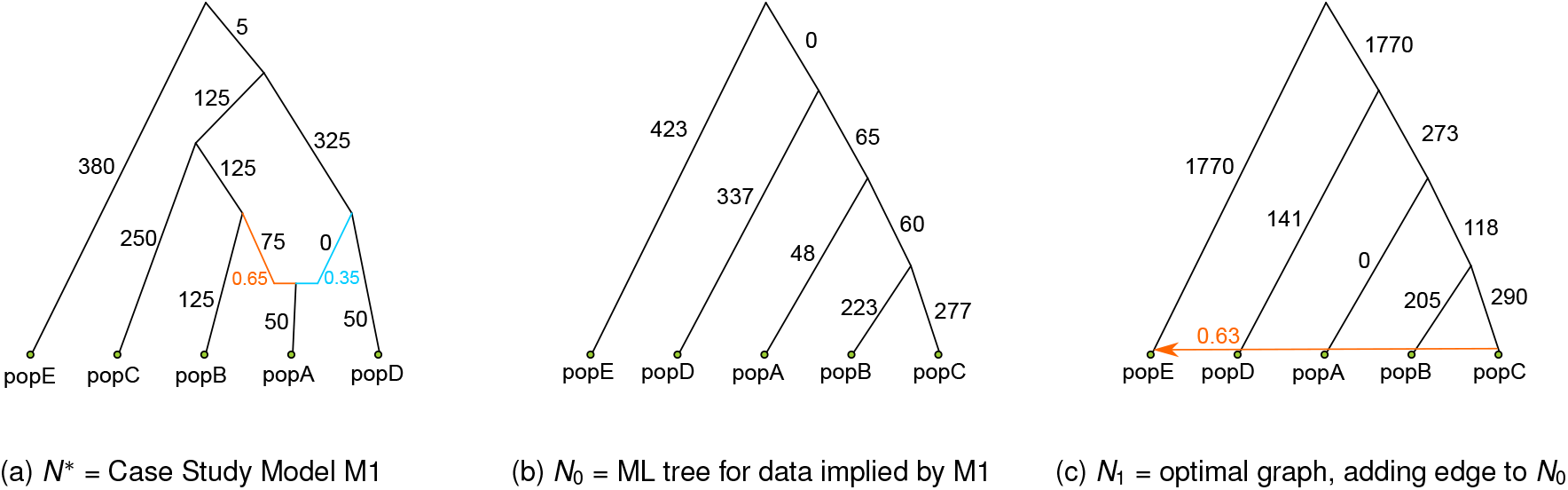
Subfigure (a) shows the model admixture graph (*N*^*\**^, **Θ**^*\**^) discussed in Section 3. We refer to this model as M1, and let ***X***^*\**^ denote the vector of *f*_2_-statistics implied by this model. Subfigure (b) shows *N*_0_: the ML (and NJ) tree for ***X***^*\**^ rooted at the outgroup *E*. Subfigure (c) shows *N*_1_: the ML network in the edge addition neighborhood of *N*_0_. Although *N*^*\**^ has the highest likelihood of all networks with one admixture node, `TreeMix`and related STB-ML methods get stuck in a local optimum, returning *N*_1_. Numerical parameters shown in subfigure (b) and (c) were computed using `TreeMix`. Log-likelihood scores, also computed using `TreeMix`, of *N*_0_, *N*_1_, and *N*^*\**^ are -1,302,625, -366,944, and 83, respectively. Branch lengths are shown multiplied by 1,000.

**Figure 2:**
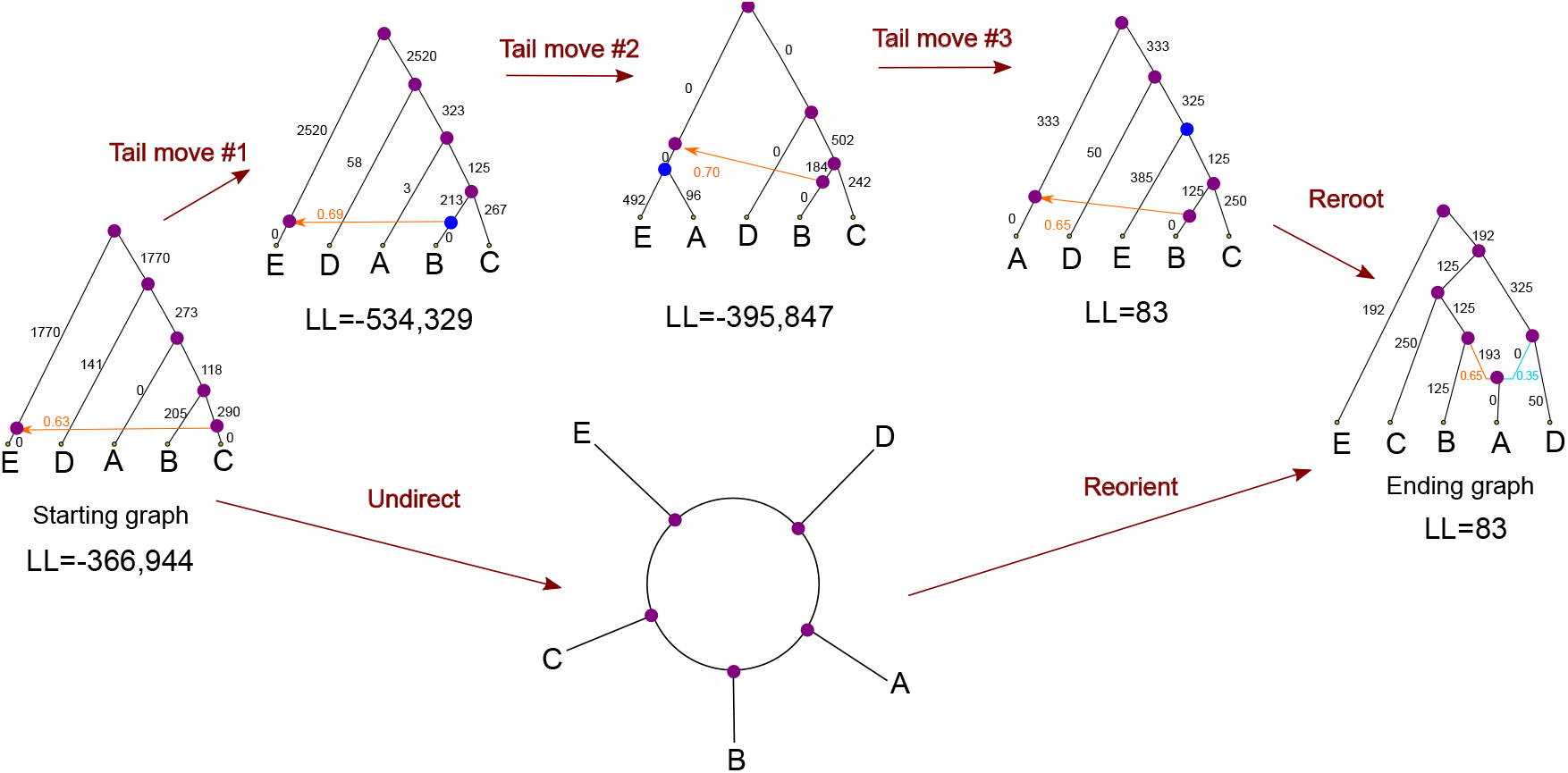
The top of the figure shows a minimal sequence of three tail moves that transforms the starting network *N*_1_ computed in Step 2 into the ending network *N*^*\**^. We found, through brute force analyses, that any sequence of tail moves from *N*_1_ to *N*^*\**^ must traverse through a network with a lower likelihood score than *N*_1_. Alternatively, *N*_1_ can be transformed into *N*^*\**^ by selecting *A* as the admixed population (instead of *E*) and redirecting the edges of the network accordingly. Adopting the language of Huber *et al*. (2019), we refer to this as reorientating the network. This is not to be confused with rerooting the network, which simply relocates the position of the root and does not change which populations are admixed. Numerical parameters and log-likelihood scores shown above were computed using `TreeMix`. Branch lengths shown are multiplied by 1,000.

#### Definition 2

(Edge Addition). An *edge addition* between two edges *e* and *f* in a directed phylogenetic network *N* involves subdividing them with new vertices *s* and *t*, respectively, and then adding an edge, referred to as the linking arc, from *s* to *t*.

#### Definition 3

(Tail Move). A *tail move* between two edges *e* and *f* in a directed phylogenetic network *N* involves relocating the tail (source vertex) of edge *e* so that it subdivides edge *f*.

Many methods require networks to have specific properties; `TreeMix`, in particular, searches for *tree-based* networks, a class of networks introduced by Francis and Steel, 2015.

#### Definition 4

(Tree-based). We say that a directed phylogenetic network *N* is *tree-based* if it can be constructed from a phylogenetic tree *T*, referred to as the base tree, via a sequence of edge additions, with the linking arcs representing gene flow, that can be performed in any order.

As shown by Francis and Steel (2015), this implies the existence of labeling *ψ* : *E*(*N*) *→ {*0, 1*}* with the following properties: first, any non-root vertex has *exactly* one of its incoming edges labeled 0, second, any internal vertex has *at least* one of its outgoing edges labeled 0, and third, the root has both of its outgoing edges labeled 0 (note this last requirement is specific to `TreeMix`). We say that edges labeled 0 are “part of the base tree,” and edges labeled 1 are “gene flow.” Labelings with these properties are called tree-based (note that there can be multiple tree-based labelings of *N*).

We conclude this section by discussing *undirected* phylogenetic networks. A phylogenetic network *N*′ = (*n*′, *S, θ*) is undirected if *n*′ is undirected and connected, with no parallel edges or self-loops (Gambette *et al*., 2012). Again, we assume that *N*′ is binary, so leaves are vertices with degree 1 and all other vertices have degree 3. It is easy to transform a directed phylogenetic network *N* into an undirected network *N*′ simply by ignoring edge directions and suppressing the vertex previously designated as the root (Gambette *et al*., 2012).

#### Definition 5

(Network Orientation). We say that a directed network *N* is an *orientation* of an undirected network *N*′ if the undirected version of *N*, denoted *N* |_*u*_, is isomorphic to *N*′.

Note that two undirected networks 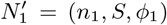 and 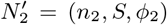 are *isomorphic* if there exists a bijection 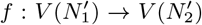 such that 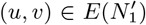 if and only if 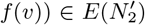 and *ϕ*_1_(*u*) = *ϕ*_2_(*f* (*u*)) for all *u*∈ *L*(*N*).

The reverse operation of transforming an undirected network *N*′ into a directed network *N*, referred to as orienting the network, is not so straightforward, because the location of the root does not uniquely determine the direction of all edges in an unrooted network (Gambette *et al*., 2012; Huber *et al*., 2019). Notably, Huber *et al*. (2019) recently showed that specifying the the location of the root (i.e. the edge that the root subdivides) and the admixture nodes in an undirected phylogenetic network *N*′ results in a unique orientation of *N*, provided that an orientation exists (Theorem 2 in Huber *et al*., 2019). Furthermore, given this information, this orientation can be can be found in linear time in the number of edges (Algorithm 1 in Huber *et al*., 2019).

In the remainder of this paper, all phylogenetic networks are directed, unless otherwise noted, and the term “phylogenetic” is often omitted when referring to phylogenetic trees and networks.

### 2.2 Likelihood Function

A phylogenetic network *N* is just one parameter in an admixture graph (*N*, **Θ**). In order to search network space, we must define a procedures for estimating numerical parameters **Θ** (i.e. the branch lengths in drift units and admixture proportions) and computing the likelihood of the resulting model given the input data.

Without loss of generality, we can define the likelihood function used by `TreeMix` (and related methods) in terms of *f*_2_-statistics. Given a model admixture graph (*N*, **Θ**), we can compute the expected value of the *f*_2_-statistics directly, up to a scaling factor (Pickrell and Pritchard, 2012; Patterson *et al*., 2012; Peter, 2016). If *N* is a tree, then the *f*_2_-statistic for populations *i, j* ∈ *S* is the sum of the lengths (in units of genetic drift) of edges on the path from the leaf labeled *i* to the leaf labeled *j* in *N*. If there is admixture, then this formula is more complex, as we must consider all paths between *i* and *j* and use the admixture proportions to weight these paths appropriately.

In practice, (*N*, **Θ**) are unknown, and each *f*_2_-statistic is estimated from allele frequency data, along its standard error. We let ***X*** and ***Z*** denote the vector of observed *f*-statistics and the variances, respectively. Given our input ***X*** and a network *N*, we can optimize **Θ** to minimize the difference between the observed *f*_2_-statistics ***X*** and the expected value of the *f*_2_-statistics for (*N*, **Θ**), denoted ***Y***. Assuming *X*_*i*_ is normally distributed with mean *Y*_*i*_ and variance *Z*_*i*_, the composite likelihood function is

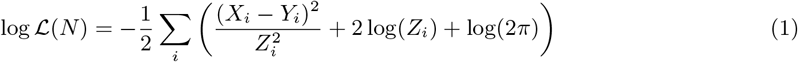

This (approximate) log-likelihood function used by `TreeMix` is related to the log-likelihood score used by `qpGraph` and `miqograph`.

As discussed by Pickrell and Pritchard (2012) and others (*e*.*g*., Lipson, 2020), the three branch lengths incident to an admixture node cannot be estimated simultaneously. Automated methods typically estimate (i.e. fit) exactly one of the three branch lengths, for example `admixturegraph` estimates the length of the outgoing edge, setting the lengths of the two (incoming) admixture edges to 0. In contrast, `TreeMix` sets the length of the outgoing edge and one of the two incoming admixture edges to 0 (note that `TreeMix` uses a tree-based labeling *ψ* of *N* to determine which admixture edges are set to 0 and which are estimated). Lastly, we note that the position of the root is not identifiable, except that it must be ancestral to admixture nodes.

## 3 Pitfalls of Starting-Tree-based ML Methods

In this section, we show that there exist simple demographic models for which STB-ML methods using the likelihood function given Equation 1 are guaranteed to get trapped in a local optimum and thus return an incorrect network.

Our case study model (*M*1), shown in Figure 1a, has five populations at the leaves, one of which is admixed. The network topology can be created by forming a caterpillar tree with leaves alphabetically labeled by the set *S* = *{A, B, C, D, E}*, rooting the tree at population *E*, and then performing an edge addition (Definition 2) on the edges incident to *D* and *A* so that *A* is admixed. In the remainder of this section, we use (*N*^*\**^, **Θ**^*\**^) to denote the network topology and numerical parameters for model *M*1.

Given (*N*^*\**^, **Θ**^*\**^), it is possible write down the *f*_2_-statistics implied by this model, denoted ***X***^*\**^. As the amount of data generated under (*N*^*\**^, **Θ**^*\**^) goes to infinity, the estimated *f*_2_-statistics converge to ***X***^*\**^, up to a scaling factor, and the standard error for every *f*_2_-statistic goes to zero (note that it is typically shifted by a small constant to avoid numerical issues).

Evaluating the likelihood of a candidate network N requires optimizing the numerical parameters **Θ**. To verify that our results were agnostic to the choices in parameter estimation by performing computational experiments using `admixturegraph` and `TreeMix`. Recall that `admixturegraph` optimizes the lengths of tree edges only, whereas `TreeMix` uses a tree-based labeling *ψ*, optimizing the lengths of the edges that are part of the base tree and whose tails (source vertices) are not admixture nodes (note that we evaluated likelihood using all tree-based labelings for a network). Scripts used in this study are available on Github: https://github.com/ekmolloy/mlno-study.

We begin our computational study by addressing whether the network topology *N*^*\**^ has the highest likelihood of all phylogenetic networks on *S* with one admixture event. We found that *N*^*\**^ has the highest likelihood through brute force, using `all graphs` function in `admixturegraph` to generate all such topologies and computing their likelihoods with both `admixturegraph` and `TreeMix`.

We now turn to the performance of STB-ML methods given ***X***^*\**^ as input. Because *M* 1 contains a single admixture event, a STB-ML method would operate by (1) searching for a ML starting tree *N*_0_, (2) searching for a ML network *N*_1_ in the edge addition neighborhood of *N*_0_, and (3) searching from *N*_1_ for a network with a higher likelihood by applying other search moves. Below, we use brute force to determine the outcome of these three steps.

**Step 1:** The first step of a STB-ML method is to estimate a starting tree on the full set of populations; this is typically done using a ML heuristic or the Neighbor Joining (NJ) algorithm (Saitou and Nei, 1987). In our case study, the NJ tree for ***X***^*\**^ is the same as the ML tree for ***X***^*\**^, which we found using brute force as described above. The NJ/ML tree rooted at outgroup *E*, denoted *N*_0_ and shown in Figure 1b, is *not* a base tree of *N*^*\**^. Note that `TreeMix` uses a random population heuristic to build a starting tree and may not be successful in finding the optimal tree in practice.

**Step 2:** The second step of a STB-ML method is to search for a ML network in the edge addition neighborhood of *N*_0_, which has size *O*(|*E*(*N*_0_)|^2^) (Definition 2). We used brute force to find the optimal network topology in this neighborhood. The ML network, denoted *N*_1_ and shown in Figure 1c, has *E* as the admixed population (note that we confirmed this outcome for all possible rootings of *N*_0_)

**Step 3:** The third step of an STB-ML method is to search from *N*_1_ for a network with a higher likelihood via hill climbing. In our brute force evaluation of all networks with one admixture node, we found that *N*_1_ has the second highest likelihood score, second only to *N*^*\**^; therefore, it is sufficient to show that *N*_1_ cannot be transformed into *N*^*\**^ via a single search move. We consider search moves that are commonly implemented in STB-ML methods, namely tail moves (Definition 3) and head moves. A head move relocates the head (target vertex) of an admixture edge to some other edge in the network, so we can verify by inspection that *N*_1_ cannot be transformed into *N*^*\**^ with head moves. To evaluate whether this can be achieved with tail moves, we implemented a subroutine that given a network *N* finds all networks in its tail move neighborhood. Applying this subroutine iteratively, we found that at least three tail moves are required to transform *N*_1_ into *N*^*\**^, modulo the position of the root, which as previously mentioned does not impact the likelihood score (note that we confirmed this outcome for all possible rootings of *N*_1_). As expected from our brute force analysis, this sequence of tail moves, shown in Figure 2, requires moving from *N*_1_ to graphs of lower likelihood, so it is impossible to reach *N*^*\**^ from *N*_1_ via hill climbing with head moves and tail moves. We conclude that our STB-ML method returns

## 4 The Maximum Likelihood Network Orientation (MLNO) Problem and OrientAGraph

In the previous section, we showed through a series of computational experiments that even for a simple model admixture graph, with just one admixed population, typical STB-ML methods can get trapped in a local optimum and return an incorrect network. In a purely hill climbing approach, we must move from incorrect network *N*_1_ to the correct network *N*^*\**^ in a single step. One possible solution is to define a search move consisting of three tail moves. The 3-tail move neighborhood of *N*_1_ has size *O*(|*E*(*N*_1_)|^6^), and it seems unlikely that a random 3-tail move would take us from *N*_1_ to *N\**. Interestingly, we can transform *N*_1_ into *N*^*\**^ simply by selecting *A* to be the admixed population and redirecting the edges of the network accordingly (Figure 2). This search strategy of considering different populations as admixed has been used by researchers (e.g. Lipson, 2020) to manually explore network space. There is a connection between this approach and network orientation, because the admixture nodes (along with a valid root position) uniquely determine the orientation of an undirected binary network (Huber *et al*., 2019). In other words, researchers manually search for a network orientation with the highest likelihood. Integrating this manual process into automated methods is appealing and leads us to propose the maximum likelihood network orientation (MLNO) problem.

### Definition 6

(Maximum Likelihood Network Orientation (MLNO) Problem). Let *N*′ be an undirected phylogenetic network, and let 𝒩(*N*′) be the orientation neighborhood of *N*′ i.e. the set of directed phylogenetic networks such that *N* ∈ 𝒩 (*N*′) implies that the undirected version of *N*, denoted *N* _*u*_, is isomorphic to *N*′. We say that a directed network *N*^*\**^ is a maximum likelihood orientation of *N*′, if *N*^*\**^ is in the set arg max_*N*∈ 𝒩(*N*′)_ *ℒ*(*N*)

Here, we take *ℒ*(*N*) to be the likelihood function given in Equation 1.

The most straightforward approach to finding the MLNO is exhaustive search, typically initiated from some directed network *N* with *h* admixture nodes. The orientation neighborhood of *N* |_*u*_ is defined by all ways of selecting *h* admixture nodes from the set *V* (*N*|_*u*_) *\ L*(*N* |_*u*_) and all ways of selecting a root edge from *E*(*N*|_*u*_) (Huber *et al*., 2019). We typically consider networks that are rooted at the outgroup *g*, denoting this set *𝒩*_*g*_ (*N*|_*u*_) ⊂ *𝒩* (*N*|_*u*_). In a brute force approach, we evaluate the likelihood of every directed network *M* ∈ *𝒩*_*g*_ (*N*|_*u*_); this requires optimizing the parameters for *M*, denoted **Θ**_*M*_.

As discussed in Sections 2.2 and 3, methods, such as `admixturegraph` and `TreeMix`, differ in how they optimize parameters, with `TreeMix` requiring a tree-based labeling *ψ* for *M*. This labeling is lost in reorientating a network, but a tree-based labeling, if one exists, can be found in linear time (Francis and Steel, 2015). An issue here is whether different tree-based labelings will yield different likelihood scores or have downstream effects on the search algorithm; however, when *h* is small, it is possible to evaluate all 2^*h*^ labelings (whether or not a given labeling is tree-based can be verified in linear time), as described by Francis and Steel (2015). We discuss this issue further in Sections 6 and 7.

In any case, an exhaustive search for a MLNO is only feasible when *h* and |*V* (*N*|_*u*_)| are sufficiently small. When *h* = 1, the orientation neighborhood *𝒩*_*g*_ (*N* |_*u*_) can be generated in *O*(|*E*(*N*|_*u*_)| *×* |*V* (*N*|_*u*_) *\ L*(*N*|_*u*_)| time, as the algorithm from Huber *et al*. (2019), which scales linearly in the |*E*(*N*|_*u*_)|, can be used to reorient *N*|_*u*_ for all ways of selecting one admixture node from the set *V* (*N*|_*u*_) *\ L*(*N*|_*u*_). For each *M* ∈ *𝒩*_*g*_ (*N*|_*u*_), we can compute the likelihood for all tree-based labelings, as there are at most two. When *h* is not fixed, we conjecture that finding the MLNO of *N*|_*u*_ is NP-hard, in which case heuristic search will be necessary.

We conclude this section by describing how we incorporate MLNO within the latest version of `TreeMix` (v1.13 Revision 231), referring to our implementation as `OrientAGraph`. To estimate an admixture graph (*N*, **Θ**) with *h* admixture events, the following steps are taken.

1. Search for a ML starting tree *N*_0_, rooting it at the outgroup.
2. For *i* = 1, 2, …, *h*:
  a. 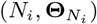 ← Search for a ML network in the (gene flow) edge addition neighborhood of *N*_*i*−1_.
  b. 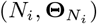 ← Search from *N*_*i*_ for a network of a higher likelihood using tails moves.
  c. 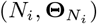 ← Search for a ML network in the (outgroup-rooted) orientation neighborhood of *N*_*i*_|_*u*_.
3. Return 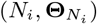

All three parts of step 2 apply operations (edge addition, tail moves, reorientations) to a network with the goal of searching for a ML network. Our case study focuses on the utility of MLNO; however, as observed in our experimental study, the relative effectiveness of these operations depends on the data set. The details of this approach are provided in the Supplementary Materials, but we summarize the differences between `TreeMix` and `OrientAGraph` below (note that Steps 1, 2b, and 3 are the same in both methods).

For step 2a, `TreeMix` searches a subset of the edge addition neighborhood, which we refer to as the gene flow edge addition neighborhood. Specifically, edge additions must occur between pairs of edges that are both labeled as part of the base tree, with the linking arc labeled as gene flow; the tree-based labeling of *N*_*i*−1_ is then extended to *N*_*i*_ in the natural way. There are still 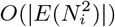 networks in this space, and `TreeMix` uses a heuristic to identify a subset of them that seem promising. In `OrientAGraph`, we enable an exhaustive search of the gene flow edge addition neighborhood. Regardless of whether a heuristic or exhaustive search is performed, the legality of edge additions still needs to be evaluated (e.g. an edge addition cannot produce cycles). Testing for legality can be sped up by performing a dynamic programming preprocessing phase prior to initializing the exhaustive search (Algorithm 1 in the Supplementary Materials). However, an exhaustive search will still be prohibitively expensive when the number of populations is large.

For Step 2c, `OrientAGraph` executes an exhaustive search for a MLNO (note this step can be performed by evaluating a single tree-based labeling found by applying the algorithm of Francis and Steel, 2015 or by evaluating all possible labelings). In the next section, we evaluate the utility of MLNO for enabling STB-ML methods using *f*-statistics to escape local optima on modestly-sized data sets. An exhaustive search for the MLNO will not scale to data sets with a large number of admixture events; we discuss how scalability might be addressed and related considerations in Section 7.

## 5 Experimental Study

In our experimental study, we utilized admixture graph models published in prior studies to benchmark `TreeMix`, `OrientAGraph`, and `miqograph`, all of which take *f*-statistics as input. The exact *f*-statistics implied by the models were given to these methods as input, representing the case of infinite data. Method performance on these model data sets can be attributed to the method itself rather than error or bias in the input. To evaluate whether the trends observed for `TreeMix` and `OrientAGraph` extended to finite data, we benchmarked these methods on genomes simulated under a subsets of the models.

### Model Data Sets

We used the `admixturegraph` R package (Leppälä *et al*., 2017) to create the *f*-statistic data sets implied by our case study model (M1) as well as seven other models (Figure 3 and Supplementary Figure S1). Model M2 is based on the toy example shown in Figure 5 of Patterson *et al*. (2012) used to benchmark `qpGraph`. All other models are based on admixture graphs estimated from biological data sets in prior studies; see Figure 3 for details. Note that M5 has the same network topology as M1; however, M1 is a toy example, whereas M5 was estimated on a biological data set by Lipson (2020). For these experiments, the standard error of the *f*-statistic was set to 0.0001 to avoid numerical problems.

**Figure 3:**
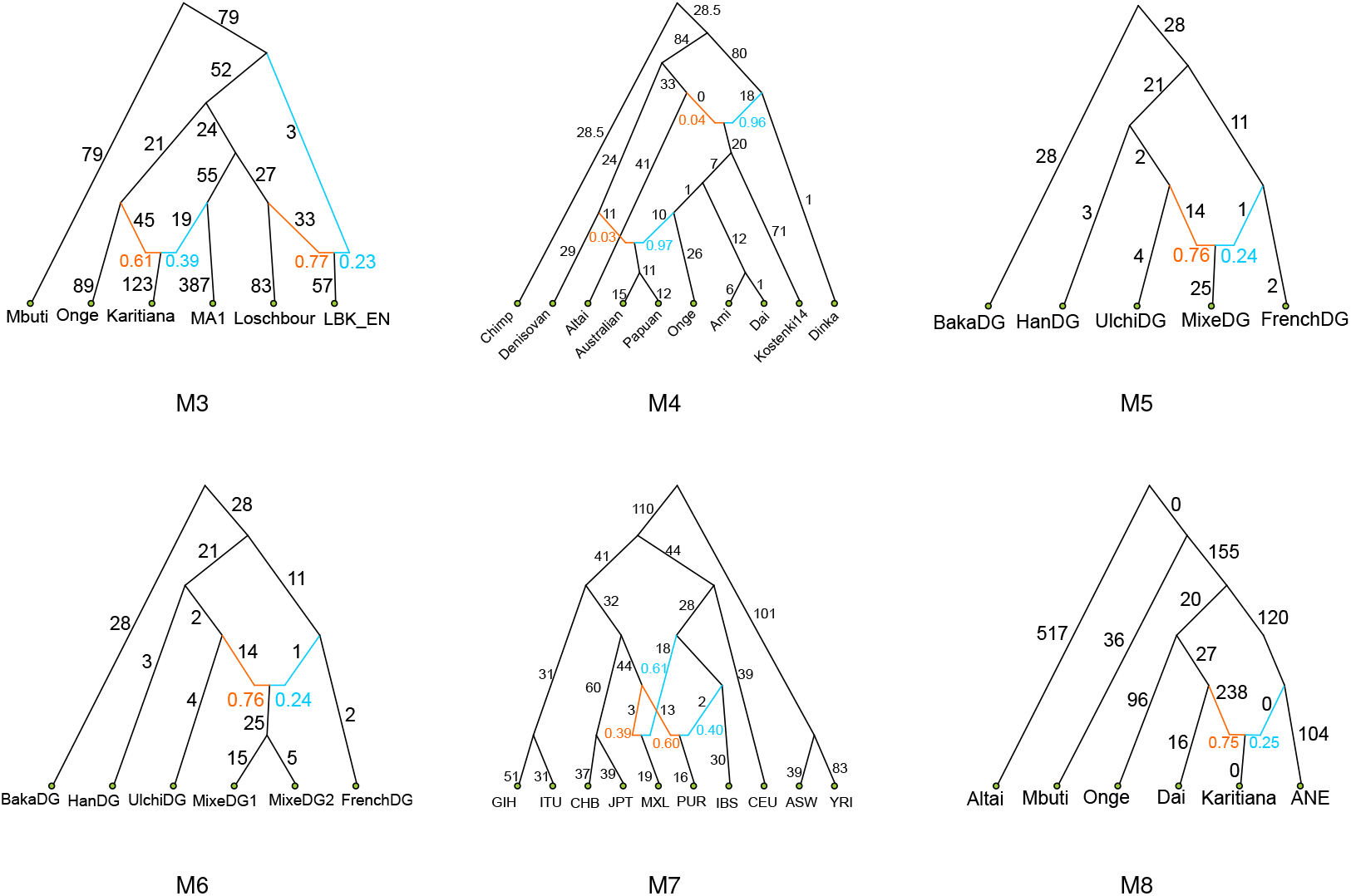
This figure shows the model demographies evaluated in our experimental study. M3 is based on Figure S8.1 from Haak *et al*. (2015), M4 is based on Figure S11.1 from Mallick *et al*. (2016) but simplified so that there are only two admixture events, M5 is based on Figure 5a from Lipson (2020), M6 is M5 but extended so that Mixe split into two populations, M7 is based on Figure 7a from Wu (2020), and M8 is based on Figure 2 from Yan *et al*. (2020). All of these admixture graphs were estimated from biological data sets. Note that the branch lengths in these subfigures are not drawn to scale and are shown multiplied by 1,000.

### Simulated Data Sets

We simulated data sets from three demographies from models (M1, M5, and M6) using `ms` (Hudson, 2002) with the number of loci set to 3, 000, the number of sites per locus set to 1 Megabase, the effective population size above the root set to 10, 000, the sample size for each population set to 20, the mutation rate set to 1.25 *×* 10^−8^, and the recombination rate set to 1 *×* 10^−8^. The models based on biological data sets had branch lengths given in units of genetic drift and were not dated, so we did not know the timing of each internal node. To simulate data with `ms`, we assigned timings at the internal nodes and then re-scaled the population size on each branch so that it had the correct length in drift units. The maximum number of SNPs per locus was approximately 6000, 3000, and 3500 for M1, M5, and M6, and we set the block size accordingly, when we used `TreeMix` to estimate *f*_2_-statistics, along with their standard error (note that we also performed experiments computing the covariance matrix instead of *f*_2_-statistics to confirm that this did not impact results). Lastly, we repeated this process of estimating matrices with `TreeMix` using SNPs from the first 100, 500, 1000, 1500, and 2000 loci. This produced a total of 18 simulated data sets.

### Method Evaluation

Our primary focus was comparing `TreeMix` and `OrientAGraph`. Recall that `OrientAGraph` is implemented on top of `TreeMix` with two different subroutines: one for exhaustive gene flow edge additions and one for MLNO (Section 4). We executed these subroutines together as well as individually to explore their relative impact. Note that we exhaustively evaluated potential tree-based labelings during the MLNO subroutine. Otherwise, methods were run with the same options: specifying the outgroup, specifying the (correct) number of admixture events, and implementing additional search moves from the starting tree, after reconstructing it with random population addition. In addition, we generated all possible population addition orders, selected 100 uniformly at random, and ran all `TreeMix` and `OrientAGraph` using each of these orders.

Unlike `TreeMix` or `OrientAGraph`, `miqograph` is guaranteed to find the true admixture graph topology *N*^*\**^, provided that *N*^*\**^ meets some topological constraints. We could not run `miqograph` on our cluster (because of how the industrial solver used by `miqograph` handles academic licenses), so we ran `miqograph` on the model data sets only. To compare running times, all analyses of model data sets were performed on the same shared computing resource (Intel(R) Xeon(R) CPU 2.10GHz server with 128 GB RAM and 32 cores). We allowed `miqograph` to use all available threads (note that `TreeMix` and `OrientAGraph` are single threaded) and gave it the same information as `TreeMix` and `OrientAGraph`, for example the outgroup and the (correct) number of admixture events. `miqograph` also requires the user to specify the granularity with which admixture proportions can be estimated (we set this value to 4) and the depth of its “search tree” (we set this value to half the number of leaves in a given model).

Methods were compared in terms of three measures: log-likelihood score of the estimated admixture graph (for `TreeMix` and `OrientAGraph` only), topological accuracy, specifically triplet distance between the true and estimated admixture graph topology (Jansson *et al*., 2019), and runtime (in seconds). For `TreeMix` and `OrientAGraph` only, we also plotted the estimated admixture graphs (for one population order) and its residuals for each model data set (see Supplementary Materials). This was done to verify that different likelihood scores corresponded to different residuals and that a triplet distance of 0 corresponded to the true admixture graph being returned.

## 6 Results and Discussion

### 6.1 Model Data Sets

In this section, we report the results of running `TreeMix`, `OrientAGraph`, and `miqograph` on model data sets (Table 1).

**Table 1:** Results for estimating admixture graphs given the *f*-statistics implied by the models in Figure 3 using `TreeMix`, `OrientAGraph` (`OAG`), and `miqograph` (`miqo`). We report the running time in seconds, the log-likelihood score (for `TreeMix` and `OrientAGraph` methods only), and the topological error (triplet distance) between the model admixture graph and the estimated admixture graph (we also note whether the correct admixture graph topology was returned). Running time is averaged across 100 runs for `TreeMix` and `OrientAGraph`.^*¶*^Indicates there was a difference in the log-likelihood score or triplet distance for the graphs returned on different population orders, in which case, we report the average.^*\**^Indicates that `miqograph` cannot get the topology correct, because of the position of the admixture event(s).^*†*^Indicates that `miqograph` did not complete within the time limit of 10,000 seconds (2.78 hours) and thus was not solved to optimality.

| Model | Log-likelihood |  | Correct Topology? |  |  | Topological Error |  |  | Running Time (secs) |  |  |
| --- | --- | --- | --- | --- | --- | --- | --- | --- | --- | --- | --- |
|  | **TreeMix** | OAG | **TreeMix** | OAG | miqo | **TreeMix** | OAG | miqo | **TreeMix** | OAG | miqo |
| M1 | -366944 | 83 | No | Yes | Yes | 9 | 0 | 0 | 0.06 | 0.19 | 398 |
| M2 | 83 | 83 | Yes | Yes | Yes | 0 | 0 | 0 | 0.04 | 0.15 | 802 |
| M3 | 124 | 124 | Yes | Yes | Yes | 0 | 0 | 0 | 0.28 | 2.12 | 2340 |
| M4* | 103¶ | 373 | No | Yes | No | 14 | 0 | 81† | 1.28 | 11.72 | 10000 |
| M5 | -33 | 83 | No | Yes | Yes | 9 | 0 | 0 | 0.05 | 0.16 | 170 |
| M6* | -30 | 124 | No | Yes | No | 15 | 0 | 2 | 0.11 | 0.31 | 556 |
| M7 | -2883 | -2883 | No | No | No | 18.6¶ | 20 | 82† | 5.58 | 13.20 | 10000 |
| M8 | 124 | 124 | Yes | Yes | Yes | 0 | 0 | 0 | 0.09 | 0.34 | 264 |

#### Models M2, M3, and M8

For these three model data sets, all methods recovered the true admixture graph topology and did so quickly.

#### Model M1

For our case study model data set, we observed the results expected based on Section 3. `TreeMix` recovered an incorrect topology, whereas `OrientAGraph` recovered the true topology. `miqograph` also recovered the true topology but was two orders of magnitude slower than `OrientAGraph`.

#### Model M4

Model M4 is an admixture graph presented with the publication of Simmons Genome Diversity data set (Mallick *et al*., 2016); however, we simplified the published model so that it only had two admixture events: gene flow from Denisova into the ancestors of Australians and Papuans as well as gene flow from Neanderthals into modern human, non-Africans. After 2.78 hours, the graph returned by `miqograph` graph had a triplet distance of 81 to the true topology. It is unclear how much this score would improve with a longer running time, given that both admixture events in M7 are not incident to the leaves, so `miqograph` is guaranteed to return an incorrect topology. In contrast, `TreeMix` and `OrientAGraph` completed in less than 15 seconds, with a triplet distance of 15 and 0 to the correct topology respectively. This difference was also reflected in the log-likelihood scores of the returned graph. These results were driven by the exhaustive search for a ML gene flow edge addition rather than a MLNO, suggesting that heuristic employed by `TreeMix` to identify candidate edge additions was ineffective for M4.

#### Models M5 and M6

On model data sets for M5 and M6, we observed similar trends to our case study (M1), that is, `TreeMix` returned an incorrect topology, which had a lower log-likelihood score than the true topology. Both `OrientAGraph` and `miqograph` returned the true topology, but again `OrientAGraph` was faster than `miqograph`.

Model M5 was estimated from Simmons Genome Diversity Project data (Mallick *et al*., 2016) by Lipson (2020), who explored the space of admixture graphs manually using `qpGraph`. We were not aware of M5 when we designed our case study, but after finding this result, we created model M6 by extending M5 to have two populations descending from the admixed population (Mixe). This enabled us to evaluate whether the trends observed were unique to the depth of the admixed population; this was not the case as the trends observed for M6 were the same as for M5. However, `miqograph` is guaranteed to recover an incorrect topology for M6 (because of its topological constraints); this was reflected in our results.

#### Summary of Models M1, M5, and M6

We confirmed that the trends observed for these three model data sets were driven by MLNO (and not an exhaustive search for a ML gene flow edge addition) by running these subroutines separately (Supplementary Table 1). We also performed exploratory analyses on these model data sets. For each model (*N*^*\**^, **Θ**^*\**^) and input data ***X***^*\**^ pair, we ran `OrientAGraph` given a base tree for *N*^*\**^ as its starting tree and using exhaustive search for ML gene flow edge additions. In this setting, `OrientAGraph` should recover the true admixture graph topology even without the MLNO; we confirmed that this was the case. We also computed the NJ tree for ***X***^*\**^ to check that it was *not* a base tree of *N*^*\**^; again, this was the case, so `TreeMix` could not be improved simply by taking the NJ tree as its starting tree.

Lastly, for each model, we scored the true admixture graph topology *N*^*\**^, using all tree-based labelings. While the parameters around the admixture node differed (as expected), the likelihood scores were very similar. While some model parameters were not identifiable by `TreeMix`, the admixed population was identifiable. Specifically, in the orientation neighborhood of *N*^*\**^, we found that `OrientAGraph` (which returned the true topology) and `TreeMix` (which returned an incorrect topology) achieved the highest and second highest likelihood scores, respectively. Importantly, the residuals showed that the true topology was a better fit to the data (Supplementary Materials Figure S3, S8, S10). Note that these results also suggest that a single tree-based labeling could be used during MLNO, at least for simple models.

#### Model M7

Model M7 was estimated by Wu (2020), who ran `GTMix` given from data from the third phase of the 1000 Genomes project (The 1000 Genomes Project Consortium, 2015). Wu (2020) found that running `TreeMix` on a related data set produced a different admixture graph topology than `GTMix`. However, `GTMix` and `TreeMix` use different inputs (gene genealogies versus *f*-statistics), different likelihood functions, and different search heuristics (although both are STB-ML methods). Our question is whether `TreeMix`’s performance could be related to its search procedure.

In principle, `TreeMix` can recover the true topology, as model M7 is tree-based; however, both `TreeMix` and `OrientAGraph` failed to recover the true admixture graph topology on this model data set, returning graphs with a log-likelihood score of -2883 for all 100 different population orders. The triplet distance between the estimated graphs to the true graph varied (either 16 or 20). We scored the true admixture graph topology given the model data set. This yielded a log-likelihood score of 373 (which is higher than -2883), so `TreeMix` is getting stuck local optimum (note that MLNO is ineffective in this scenario).

`miqograph` also fails, returning a topology with a triplet distance of 82 from the correct topology. Although `miqograph` can recover the correct graph as both admixture nodes are incident to leaves, it did not solve its problem to optimality within our allowed time frame of 10,000 seconds. This suggests that users may want to be wary of the results produced by `miqograph`, when it does not succeed in solving its problem to optimality. Interestingly, M7 is the only model that we studied that is where a single vertex is a source for two different admixture edges (i.e. it is tree-child); this model may be of particular interest to method developers.

#### Running time comparisons

The difference in running time between `OrientAGraph` and `TreeMix` was not pronounced on the six model data sets with at most 7 populations at the leaves and 2 admixture events (see Table 1). On these data sets, `miqograph` was slower than either `OrientAGraph` and `TreeMix` but still solved its problem to optimality. `miqograph` did not solve its problem to optimality within the allowed time frame for our study (10,000 seconds) on the remaining two model data sets, with 10 populations and 2 admixture events. On these data sets, there was noticeable slow down in the running time of `OrientAGraph` compared to `TreeMix`. These results are expected as `OrientAGraph` is the same as `TreeMix` but does more work; we discuss the issue of scalability in Section 7.

### 6.2 Simulated Data Sets

We used genome-scale data sets simulated from models M1, M5, and M6 to evaluate whether the trends observed for `TreeMix` and `OrientAGraph` on model data sets (representing infinite data) extended to the case of finite data (Figure 4). This was the case: `OrientAGraph` recovered the true topology, and `TreeMix` recovered an incorrect topology, on simulated data sets with *>* 1,000 loci (Figure 4). For M5 and M6, both `TreeMix` and `OrientAGraph` sometimes failed to recover the true topology for smaller numbers of loci (≤ 1,000 loci), likely due to differences between the estimated *f*_2_-statistics and the expected *f*_2_-statistics for the true admixture graph.

**Figure 4:**
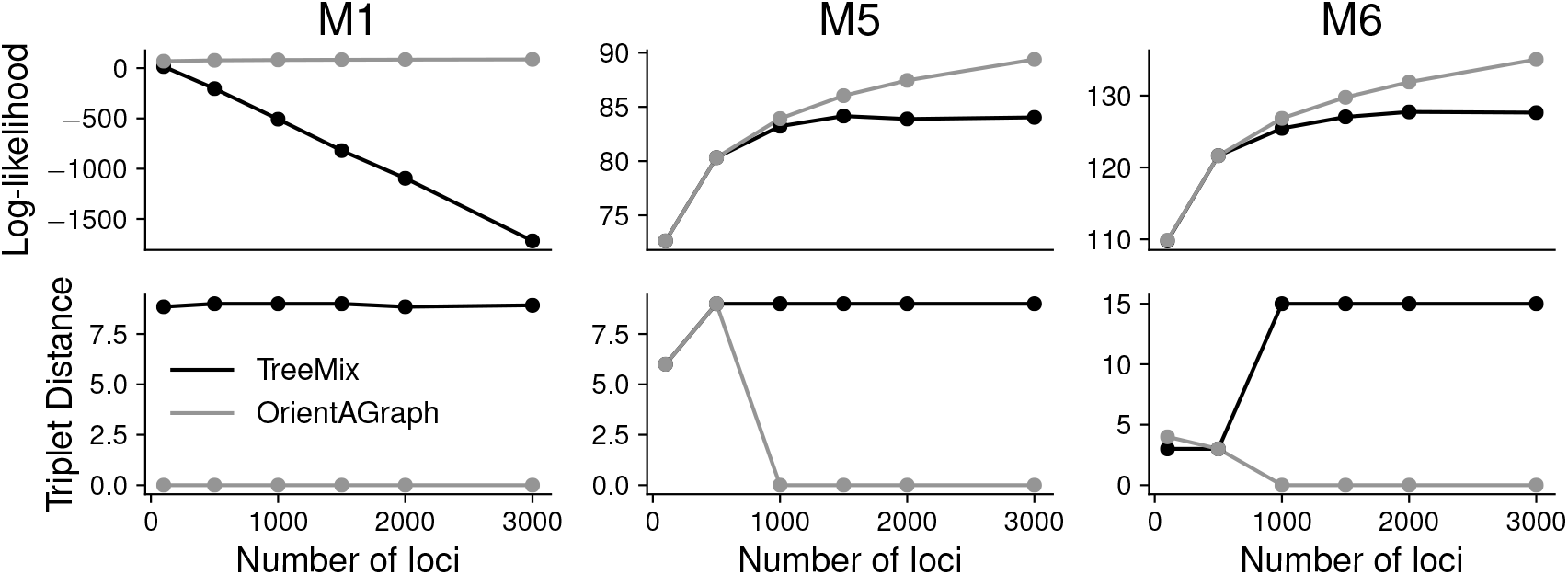
Results from simulated data ranging from 100 loci to 3,000 loci. Note that we are showing averages across runs of `TreeMix` for each population order (120 combinations for M1 and M5 and 720 combinations for M6). On these data sets, the results are consistent across runs so we only show the average. While we show both log-likelihood and triplet distance, we do not expect these measures to be fully correlated (i.e. lower triplet score may not imply lower log-likelihood).

## 7 Conclusions and Future Work

In this work, we proposed a new search strategy based on network orientation and explored its utility in the context of starting-tree-based maximum likelihood (STB-ML) methods that take *f*-statistics as input. Our current implementation, referred to as `OrientAGraph`, relies on an exhaustive search for a maximum likelihood network orientation (MLNO). This is compounded by finding a tree-based labeling, as `OrientAGraph` is implemented on top of `TreeMix`, which requires such a labeling to evaluate its likelihood function. This points to the broader challenge of fitting numerical parameters from *f*-statistics in an automated fashion. Scalability might be improved by implementing a procedure for fitting model parameters that does not require a base tree (e.g. `admixturegraph`). However, this does not address the fact that there are inherent limitations in terms of parameter identifiability. As shown by Lipson (2020), it is possible for two admixture graphs with different topologies to fit the observed *f*-statistics perfectly and thus have the same likelihood score (Equation 1). We worked to mitigate the issue of parameter identifiability in our case study and computational experiments in two ways: first, we evaluated whether differences in topological accuracy corresponded to differences in likelihood scores and residuals, and second, we selected model admixture graphs based on the topological considerations discussed by Lipson (2020).

Beyond the likelihood function, method performance can be impacted by error or bias in the input data and the search heuristic. In this work, we focused on the search heuristic by considering the case of infinite data (as well as finite data) and by performing some additional experiments. For example, we found that MLNO was unnecessary to recover true admixture graph topology *N*^*\**^ when `TreeMix` was given a base tree of *N*^*\**^ as its starting tree. We also confirmed that the NJ tree for the *f*_2_-statistics implied by the true admixture graph was not a base tree of *N*^*\**^, so the issue is not specific to searching for a ML starting tree. One possibility is that methods based *f*_2_-statistics are particularly susceptible to bad starting trees given the information content of their input. This could explain why a recent study by Cao *et al*. (2019) found an STB-ML method that takes estimated genealogical trees as input to be relatively accurate. Genealogical trees have more information content, especially when rooted; however, estimating them accurately is challenging from both a computational and statistical perspective. Furthermore, likelihood functions based genealogical trees are far more computationally intensive than those based on *f*-statistics. In any case, exploring the utility of MLNO for ML methods that use different inputs and/or Bayesian methods that sample (rather than hill climb) network space are interesting directions for future research.

Another important direction is scalability. `OrientAGraph`, while efficient for the admixture graphs considered here, will not scale to large numbers of populations and/or admixture events. In this case, we need to constrain the search for a MLNO, perhaps by maintaining a set of nodes that are required to be admixed across iterations. For example, in iteration *i*, our search of the orientation neighborhood of *N*_*i*_ can be constrained to *O*(|*V* (*N*_*i*_)|) networks, if the subset of nodes that are admixed in network *N*_*i*−1_ are required to be admixed in network *N*_*i*_. This amounts to reorientating network *N*_*i*_ “around” the *i*^*th*^ edge addition only. Such a heuristic seems promising in the scenario that admixture events are sufficiently decoupled (e.g. the network is level-1) but may not be effective otherwise. Broadly speaking, developing fully automated inference methods that are accurate yet scalable under complex admixture scenarios is an important direction for future work.

## Acknowledgments and Funding

We thank the four anonymous reviewers for their comments that led to improvements in this work. This work was funded by the following grants: NSF GRFP DGE-1650604 (A.D.), NIH R35GM125055 (S.S. and E.K.M.), and an Alfred P. Sloan Research Fellowship (S.S.).

## References

Cao, Z., Zhu, J., and Nakhleh, L. (2019). Empirical performance of tree-based inference of phylogenetic networks. In K. T. Huber and D. Gusfield, editors, 19th International Workshop on Algorithms in Bioinformatics, WABI 2019, September 8-10, 2019, Niagara Falls, NY, USA, volume 143 of LIPIcs, pages 21:1–21:13. Schloss Dagstuhl - Leibniz-Zentrum für Informatik.

Edelman, N. B., Frandsen, P. B., Miyagi, M., et al. (2019). Genomic architecture and introgression shape a butterfly radiation. Science, 366(6465), 594–599.

Francis, A. R. and Steel, M. (2015). Which Phylogenetic Networks are Merely Trees with Additional Arcs? Systematic Biology, 64(5), 768–777.

Gambette, P., Berry, V., and Paul, C. (2012). Quartets and Unrooted Phylogenetic Networks. Journal of Bioinformatics and Computational Biology, 10(04), 1250004.

Gambette, P., van Iersel, L., Jones, M., Lafond, M., Pardi, F., and Scornavacca, C. (2017). Rearrangement moves on rooted phylogenetic networks. PLOS Computational Biology, 13(8), 1–21.

Green, R. E., Krause, J., Briggs, A. W., et al. (2010). A Draft Sequence of the Neandertal Genome. Science, 328(5979), 710–722.

Haak, W., Lazaridis, I., Patterson, N., et al. (2015). Massive migration from the steppe was a source for Indo-European languages in Europe. Nature, 522, 207–211.

Harney, E., Patterson, N., Reich, D., and Wakeley, J. (2021). Assessing the performance of qpAdm: a statistical tool for studying population admixture. Genetics, 217, iyaa045.

Huber, K. T., van Iersel, L., Janssen, R., Jones, M., Moulton, V., Murakami, Y., and Semple, C. (2019). Rooting for phylogenetic networks. arXiv, CoRR, abs/1906.07430.

Hudson, R. R. (2002). Generating samples under a Wright–Fisher neutral model of genetic variation. Bioinformatics, 18(2), 337–338.

Janssen, R., Jones, M., Erdos, P., van Iersel, L., and Scornavacca, C. (2018). Exploring the Tiers of Rooted Phylogenetic Network Space Using Tail Moves. Bulletin of Mathematical Biology, 80(8), 2177–2208.

Jansson, J., Mampentzidis, K., Rajaby, R., and Sung, W.-K. (2019). Computing the Rooted Triplet Distance Between Phylogenetic Networks. In C. J. Colbourn, R. Grossi, and N. Pisanti, editors, Combinatorial Algorithms, pages 290–303, Cham. Springer International Publishing.

Leppälä, K., Nielsen, S. V., and Mailund, T. (2017). admixturegraph: an R package for admixture graph manipulation and fitting. Bioinformatics, 33(11), 1738–1740.

Lipson, M. (2020). Applying f4-statistics and admixture graphs: Theory and examples. Molecular Ecology Resources, 20(6), 1658–1667.

Lipson, M., Loh, P.-R., Levin, A., Reich, D., Patterson, N., and Berger, B. (2013). Efficient Moment-Based Inference of Admixture Parameters and Sources of Gene Flow. Molecular Biology and Evolution, 30(8), 1788–1802.

Lipson, M., Loh, P.-R., Patterson, N., Moorjani, P., Ko, Y.-C., Stoneking, M., Berger, B., and Reich, D. (2014). Reconstructing Austronesian population history in Island Southeast Asia. Nature Communications, 5(1), 4689.

Mallick, S., Li, H., Lipson, M., et al. (2016). The Simons Genome Diversity Project: 300 genomes from 142 diverse populations. Nature, 538, 201—-206.

McDiarmid, C., Semple, C., and Welsh, D. (2015). Counting Phylogenetic Networks. Annals of Combinatorics, 19, 205–224.

Patterson, N., Moorjani, P., Luo, Y., Mallick, S., Rohland, N., Zhan, Y., Genschoreck, T., Webster, T., and Reich, D. (2012). Ancient Admixture in Human History. Genetics, 192(3), 1065–1093.

Peter, B. M. (2016). Admixture, Population Structure, and F-Statistics. Genetics, 202(4), 1485–1501.

Pickrell, J. K. and Pritchard, J. K. (2012). Inference of Population Splits and Mixtures from Genome-Wide Allele Frequency Data. PLOS Genetics, 8(11), 1–17.

Pilot, M., Moura, A., Okhlopkov, I., et al. (2019). Global Phylogeographic and Admixture Patterns in Grey Wolves and Genetic Legacy of An Ancient Siberian Lineage. Scientific Reports, 9, 17328.

Saitou, N. and Nei, M. (1987). The neighbor-joining method: a new method for reconstructing phylogenetic trees. Molecular Biology and Evolution, 4(4), 406–425.

The 1000 Genomes Project Consortium (2015). A global reference for human genetic variation. Nature, 526, 68–74.

Wen, D., Yu, Y., Zhu, J., and Nakhleh, L. (2018). Inferring Phylogenetic Networks Using PhyloNet. Systematic Biology, 67(4), 735–740.

Wu, Y. (2020). Inference of population admixture network from local gene genealogies: a coalescent-based maximum likelihood approach. Bioinformatics, 36(Supplement 1), i326–i334.

Yan, J., Patterson, N., and Narasimhan, V. (2020). miqoGraph : Fitting admixture graphs using mixed-integer quadratic optimization. Bioinformatics. btaa988.

Yu, Y., Dong, J., Liu, K. J., and Nakhleh, L. (2014). Maximum likelihood inference of reticulate evolutionary histories. Proceedings of the National Academy of Sciences, 111(46), 16448–16453.

