## Supplement for "Advancing admixture graph estimation via maximum likelihood network orientation"

May 23, 2021

### 1 Supplementary Methods

#### 1.1 Overview

In Section 4 of the main paper, we provide a high-level algorithm description for `OrientAGraph`, which takes a vector $\boldsymbol{X}$ of $f$-statistics as input and searches for a maximum likelihood (ML) admixture graph $(N, \boldsymbol{\Theta}_N)$ with $h$ admixture events. The ML search proceeds as follows.

1. Search for a ML starting tree $N_0$, rooting it at outgroup $g$.
2. For $i = 1, 2, \ldots, h$:
   (a) $(N_i, \boldsymbol{\Theta}_{N_i}) \leftarrow$ Search for a ML network in the gene flow )edge addition neighborhood of $N_{i-1}$.
   (b) $(N_i, \boldsymbol{\Theta}_{N_i}) \leftarrow$ Search from $N_i$ for a network with higher likelihood using tail moves.
   (c) $(N_i, \boldsymbol{\Theta}_{N_i}) \leftarrow$ Search for a ML network in the (outgroup-rooted) orientation neighborhood of $N_i|_u$.
3. Return $(N_i, \boldsymbol{\Theta}_{N_i})$

`OrientAGraph` is implemented on top of `TreeMix` [5]. Steps 1, 2b, and 3 are the *same* as `TreeMix`. Step 2a can be performed using the same heuristic as `TreeMix` or using an exhaustive search (Sections 1.3 and 1.4); the exhaustive search can be executed by using the `-allmigs` option in `OrientAGraph`. Step 2b is new to `OrientAGraph` and can be performed using a heuristic or an exhaustive search for the MLNO (Section 1.5). The exhaustive search can be executed using the `-mlno` options in `OrientAGraph`; we plan to implement the heuristic described in Section 7 in the near future.

#### 1.2 Data Structures

The admixture graph data structure implemented by `TreeMix`, and thus `OrientAGraph`, must include a directed phylogenetic network $N^C$ with an edge labeling $\mathcal{L} : E(N^C) \to \{0, 1\}$. The network $N^C$ must have the following properties:

- the root is a vertex with indegree 0 and outdegree 2,
- the leaves are vertices with indegree $\geq 1$ and outdegree 0, and
- internal vertices have indegree $\geq 1$ and outdegree 2.

The edge labeling $\psi$ must have the following properties:

- for the root vertex, *both* of the outgoing edges must be labeled 0,
- for any non-root vertex, *exactly one* of the incoming edges must be labeled 0, and
- for any non-root, non-leaf vertex, *at least one* of the outgoing edges must be labeled 0.

Such an edge labeling $\psi$ is said to be tree-based, and we can verify whether a given an edge labeling for $N^C$ is tree-based in $O(|V(N^C)|)$ time when the number $h$ of admixture nodes is fixed. Any network $N$ for which a tree-based labeling exists is referred to as tree-based; a tree-based labeling for $N$, if one exists, can be found in linear time via a reduction to 2-SAT [1].

Although $N^C$ is not a *binary* phylogenetic network, it can be viewed as such, as we now describe. Let $N$ be a binary, directed phylogenetic network, and let $\psi$ be a tree-based labeling of $N$. Because $N$ is binary, every admixture node in $N$ has indegree 2 and outdegree 1. For some admixture node in $N$, let $e, f$ denote the two incoming edges and let $g$ denote the outgoing edge. Because $\psi$ is tree-based, either $e$ and $g$ are drawn as part of base tree (in which case $f$ is drawn as gene flow); otherwise, $f$ and $g$ are drawn as part of the base tree (in which case $e$ is drawn as gene flow). As discussed in Section 2.2 of the main paper, in the former case, `TreeMix` forces $f$ and $g$ to have length 0, and in the latter case, `TreeMix` forces $e$ and $g$ to have length 0. It follows that we do not need to store edge $g$ and can contract it (Definition 1). Contracting edges in $N$ until there are no edges with outdegree 1 produces a *contracted version* of the binary phylogenetic network $N$, denoted $N^C$, that satisfies the requirements given above. This is the data structure implemented by `TreeMix`, and thus `OrientAGraph`.

**Definition 1** (Refinements and Contractions)**.** *An* edge contraction *corresponds to deleting edge* $e = u \mapsto v$ *but not its vertices* $u$ *and* $v$ *from the graph* $G$ *and then identifying* $u$ *and* $v$ *(i.e. combining* $u$ *and* $v$ *into a single vertex in the natural way). A* vertex refinement *is simply the opposite of a contraction.*

#### 1.3 Edge Addition Neighborhood

In Step 2a, `TreeMix` and `OrientAGraph` search for a ML network in what we will call the *gene flow edge addition neighborhood* of a given the contracted version of a binary network $N_i^C$ with tree-based labeling $\psi_i$. First, we consider the *edge addition neighborhood* of a binary network. An edge addition takes a binary network $N_i$ as input and returns a binary network $N_{i+1}$ such that there exists an edge $e \in E(N_{i+1})$ whose deletion from $N_{i+1}$ produces a network that is isomorphic to $N_i$, after suppressing vertices with indegree 1 and outdegree 1. This is encoded as the following operation.

**Definition 2** (Edge Addition Operation)**.** *An* edge addition *is an operation in which two different edges of a binary, directed phylogenetic network* $N_i$*, the source edge* $e_s = u_s \mapsto w_s$ *and the target edge* $e_t = u_t \mapsto w_t$*, are subdivided with new vertices* $v_s$ *and* $v_t$*, respectively, and then an edge* $e'$ *is added from* $v_s$ *to* $v_t$*. An edge addition is only* legal *if the resulting network* $N_{i+1}$ *is a binary, directed phylogenetic network.*

Now we consider the case where $N_i$ has a tree-based labeling $\psi_i$. If we require the new edge $e'$ to be labeled as gene flow in $N_{i+1}$ and $e_s$ and $e_t$ to be labeled as part of the base tree in $N_i$, then the labeling $\psi_i$ can be extended to $N_{i+1}$, denoted $\psi_{i+1}$, in the natural way (i.e. $\psi_{i+1}(e') = 1$ and $\psi_{i+1}(u_s \mapsto v_s) = \psi_{i+1}(v_s \mapsto w_s) = 0$). Nothing is done for the remaining edges (i.e. $\psi_i(e) = \psi_{i+1}(e)$ for all $e \in E(N_i) \cap E(N_{i+1})$). We refer to edge additions of this form as being in the *gene flow edge addition neighborhood* of $N_i$.

Recall that `TreeMix` and `OrientAGraph` operate on the contracted version of a binary network, so they would immediately contract the edge $v_t \mapsto w_t$. Therefore, edge additions can be implemented on the contracted version of a binary network $N_i^C$ by subdividing an edge $e_s = u_s \mapsto w_s \in E(N_i^C)$ with a new vertex $v_s$, and then adding an edge $e'$ from $v_s$ to some target vertex $v_t \in V(N_i^C)$. In order for this edge addition to be legal, $w_s \neq v_t$ (as the edge addition would produce a parallel edge in $N_{i+1}^C$) and $v_t$ cannot be on a some directed path from the root to $w_s$ (as the edge addition would produce a cycle in $N_{i+1}^C$). `TreeMix` and `OrientAGraph` constrain their search to the gene flow edge addition neighborhood, so $e_s$ must be labeled as part of the base tree in $N_i^C$ (note that we can view the edge addition as being on the incoming edge to $v_t$ that is labeled as part of the base tree). Lastly, `TreeMix` restricts edge additions to meet some additional requirements that are likely important for preventing issues when estimating numerical parameters (see function `PopGraph::is_legal_migration`). It appears that these constraints can be verified by looking locally around $e_s$ and $v_t$. We will broadly refer to edge additions that meet the requirements of `TreeMix` (and thus `OrientAGraph`) as being in the *gene flow edge addition neighborhood* of $N_i^C$.

#### 1.4 Maximum Likelihood Edge Addition

We now turn to the task of searching for a ML network in the gene flow edge addition neighborhood of $N_i^C$, defined in Section 1.3. `TreeMix` builds a subset of possible edge additions in this neighborhood that seem promising by using the residual matrix to identify pairs of populations for which the current model is a poor fit to the input data. For each edge addition in this subset, `TreeMix` determines the legality and then, if possible, fits parameters and evaluates the log-likelihood function. The most computationally intensive step of checking the legality of an edge addition on edge $e_s = u_s \mapsto w_s$ and vertex $v_t$ is to determine whether $v_t$ is on any directed path from the root to $w_s$ (because an edge addition in this case produces a cycle). `OrientAGraph` enables an exhaustive search of the gene flow edge addition neighborhood (Algorithm 1), so it makes sense to compute this information for each vertex in a dynamic programming preprocessing step. Afterward, the legality of an edge addition is fast to compute. However, if an edge addition is legal, then we still need to fit parameters and evaluate the log-likelihood function. Doing this work for the $|V(N_i^C)| \times |E(N_i^C)|$ possible (gene flow) edge additions will not scale to large admixture graphs, although it is suitable for modestly-sized graphs.

#### 1.5 Maximum Likelihood Network Orientation

Lastly, we discuss how to find the maximum likelihood network orientation (MLNO) of a binary, directed phylogenetic network $N$ with root $r$ and $h$ admixture nodes (i.e. vertices with indegree $> 1$). Let $Y = V(N) \setminus L(N) \setminus \{r\}$, and recall that an orientation of the undirected version of $N$, denoted $N|_u$, is uniquely determined by a set $A \subseteq Y$ of $h$ admixture nodes (i.e. nodes with indegree 2) and an edge $e_r$ that determines the position of the root [2]. If a valid orientation exists for some pair $A, e_r$, it can be found in linear time by running Algorithm 2 (adapted from [2]). To exhaustively search the orientation neighborhood of $N$, we consider the directed networks found by orientating $N|_u$ for every possible subset of $Y$ with size $h$ (note that we fix the root $r$ on an edge $e_r$, typically chosen so that the root is incident to an outgroup).

There are some technicalities to consider as `OrientAGraph` operates on the contracted-version of a binary phylogenetic network, denoted $N^C$, with tree-based labeling $\psi$. First, we need to refine vertices in $N^C$ (Definition 1), until every vertex with indegree 2 has outdegree 1. If all vertices in $N^C$ have indegree $\leq 2$, then applying this process to $N^C$ produces a unique binary network. Otherwise, the resulting network, denoted $N^R$, will have stacked admixture nodes (i.e. there exists an edge that has an admixture node as its source and its target) and the branching order of admixture nodes in this stack will have been arbitrarily resolved. It is possible that the chosen branching order could impact orientation, but we speculate that if there exists a set of stacked admixture nodes, the utility of orientation will be greatest for the first edge addition that produces an admixture node in a stack (also note that there will be issues with parameter identifiability in the case of stacked admixture nodes). Then, after applying the orientation algorithm, we need form the contracted version of the resulting network and evaluate its likelihood using a tree-based labeling. The entire process is summarized in Algorithm 3.

We want to emphasize that this work is only necessary because we implemented MLNO on top of `TreeMix`, with the goal of taking advantage of `TreeMix`'s existing functions for fitting parameters, computing likelihood, etc. Moving forward, it would be advantageous, at least from the perspective of computational complexity, to take the approach of `admixturegraph` (setting all admixture edges to length 0) so that we can optimize parameters without finding a tree-based labeling (and to store a binary tree rather than the contracted version of one). For these reasons, we did not focus on optimizing the compute kernels specific to `TreeMix`.

##### 1.6 Algorithms

**Algorithm 1:** Exhaustive search ML network in the (gene flow) edge addition neighborhood of $N_i^C$; see Section 1.4 above and Section 4 in the main paper.

**Input :** A network $N_i^C$ with tree-based labeling $\psi_i$, and the vector $\boldsymbol{X}$ of observed $f$-statistics (note that $N_i^C$ is the contracted version of a binary network)

**Output:** A ML network $N_{i+1}^C$ with tree-based labeling $\psi_{i+1}$ in the gene flow edge addition neighborhood of $N_i^C$ (note that $N_{i+1}^C$ is the contracted version of a binary network). If the result is a network with a worse score than $N_i^C$, then $N_i^C$ is returned.

---

**Function** `FindMLGFEdgeAdditionExhaustive(`$N_i^C, \psi_i, \boldsymbol{X}$`)`:
  $(N_i^C, \boldsymbol{\Theta}_{N_i^C}) \leftarrow$ `Fit(`$N_i^C, \psi_i$`)`
  $s_{best} \leftarrow$ `LogLikelihood(`$N_i^C, \boldsymbol{\Theta}_{\boldsymbol{N_i^C}}, \boldsymbol{X}$`)`
  $N_{best} \leftarrow N_i^C$; $\psi_{best} \leftarrow \psi_i$
  **for** $(e_s, v_t) \in$ `GetGFEdgeAdditionNeighborhood(`$N_i^C, \psi_i$`)` **do**
    $(N_{i+1}^C, \psi_{i+1}) \leftarrow$ `AddGeneFlowEdgeToBaseTree(`$N_i^C, \psi_i, e_s, v_t$`)`
    $(N_{i+1}^C, \boldsymbol{\Theta}_{N_i^C}) \leftarrow$ `Fit(`$N_{i+1}^C, \psi_{i+1}$`)`
    $s \leftarrow$ `LogLikelihood(`$N_{i+1}^C, \boldsymbol{\Theta}_{\boldsymbol{N_{i+1}C}}, \boldsymbol{X}$`)`
    **if** $s > s_{best}$ **then**
      $N_{best} \leftarrow N_{i+1}^C$; $\psi_{best} \leftarrow \psi_{i+1}$; $s_{best} \leftarrow s$
  **return** $N_{best}, \psi_{best}, s_{best}$

**Function** `GetGFEdgeAdditionNeighborhood(`$N^C, \psi$`)`:
  $A \leftarrow \emptyset$
  $P_{root} \leftarrow$ `GetAllPathsToRoot(`$N^C$`)`
  **for** $e_s = u_s \mapsto w_s \in E(N_i^C)$ **do**
    **for** $v_t \in V(N^C)$ **do**
      **if** $\psi(e_s) = 1$ **then** continue
      **if** $w_s = v_t$ **then** continue
      **if** $v_t \in P_{root}[w_s]$ **then** continue
      `// Verify other (local) requirements of TreeMix so that edge additions do not cause numerical issues.`
      $A \leftarrow A \cup \{(e_s, v_t)\}$
  **return** $A$

**Function** `GetAllPathsToRoot(`$N^C$`)`:
  $P_{root} \leftarrow$ `Dictionary()`
  **for** $v \in$ `BreadthFirstTraversal(`$N^C$`)` **do**
    $P_{root}[v] \leftarrow \emptyset$
    **for** $p \in$ `GetParents(`$v$`)` **do** $P_{root}[v] \leftarrow P_{root}[v] \cup \{p\} \cup P_{root}[p]$
  **return** $P_{root}$

---

**Algorithm 2:** Reorient a directed phylogenetic network given a set of admixture nodes and a location for the root using the algorithm from [2].

**Input :** A directed binary phylogenetic network $N$, a set $A \subset V(N)$ of vertices to be designated as admixture nodes, and an edge $e_r \in E(N)$ on which to place the root.
**Output:** A reoriented version of $N$

---

```
Function Reorient(N, A, e_r):
    // Step 1: Set current/desired in-degrees of vertices, and set edges as
       undirected.
    r ← GetRoot(N)
    for v ∈ V(N) \ {r} do
        v.currentIndegree ← 0
        if v ∈ A then v.desiredIndegree ← 2
        else v.desiredIndegree ← 1
    for e ∈ E(N) do e.isDirected ← False

    // Step 2: Move the root vertex r to edge e_r.
    if GetSource(e_r) != r then
        {x, y} ← GetChildren(r)
        AddEdge(x ↦ y, N)
        DeleteEdge(r ↦ x, N); DeleteEdge(r ↦ y, N)
        s ← GetSource(e_r); t ← GetTarget(e_r)
        AddEdge(r ↦ s, N); AddEdge(r ↦ t, N)
        DeleteEdge(s ↦ t, N);

    // Step 3: Direct the edges.
    n ← 0; vqueue ← Queue(); vqueue.PushBack(r)
    while vqueue is not empty do
        v ← vqueue.Pop()
        X ← ∅
        for e ∈ GetOutgoingEdges(v) do
            if not e.isDirected then X ← X ∪ {e}
        for e ∈ GetIncomingEdges(v) do
            if not e.isDirected then
                e ← SwapSourceAndTarget(e)
                X ← X ∪ {e}
        for e ∈ X do
            e.isDirected ← True; n ← n + 1; t ← GetTarget(e)
            t.currentIndegree ← t.currentIndegree + 1
            if t.currentIndegree == t.desiredIndegree then vqueue.PushBack(t)

    // Step 4: Check for success.
    if n == |E(N)| then return N
    else return Null
```

**Algorithm 3:** Exhaustive search ML network in the orientation neighborhood of $N_i^C$; see Section 1.5 above and Section 4 in the main paper. Note that this is pseudocode showing the algorithm structure and that the work related to transforming data structures and evaluating different labelings could be avoided by optimizing parameters differently, i.e. it is a result of building on top of existing code.

**Input :** A network $N^C$ (with tree-based labeling $\psi$, $h$ admixture nodes, and root $r$ with edge $e_r$ incident to the outgroup), and the vector $\boldsymbol{X}$ of observed $f$-statistics (note that $N^C$ is the contracted version of a binary network)

**Output:** A MLNO network of $N^C$ with a tree-based labeling.

```
Function FindMLNOExhaustive(N^C, ψ, X):
    (N^C, Θ_{N^C}) ← Fit(N^C, ψ)
    s_best ← LogLikelihood(N^C, Θ_{N^C}, X)
    N_best ← N^C; ψ_best ← ψ
    for N^C ∈ GetOrientationNeighborhood(N^C) do
        for ψ ∈ GetBaseTreeNeighborhood(N) do
            (N^C, Θ_{N^C}) ← Fit(N^C, ψ)
            s ← LogLikelihood(N^C, Θ_{N^C}, X)
            if s > s_best then
                N_best ← N^C; ψ_best ← ψ; s_best ← s
    return N_best, ψ_best, s_best

Function GetOrientationNeighborhood(N^C):
    N ← Refine(N^C)
    Y ← V(N) \ L(N) \ {r}
    Z ← ∅
    for A ∈ AllSubsetsOfSize(Y, h) do
        M ← Reorient(N, A, e_r)
        M ← Contract(M)
        Z ← Z ∪ M
    return Z
```

#### 1.7 Topological Accuracy

We computed topological accuracy as the triplet distance compute the true and estimated admixture graph topology, using the first algorithm from [3] implemented in `ntd`. Triplets are rooted phylogenetic trees on three leaves and can be identified in a phylogenetic network by restricting it to three populations; if at least one of the populations is admixed, the restricted graph implies multiple triplets. The triplet distance between two networks is the number of triplets in the true graph that are not in the estimated graph plus the number of triplets in the estimated graph that are not in the true graph. Before computing the triplet distance, we rooted all estimated networks at the outgroup—this changes the position of the root but NOT the admixed populations. In some cases, `TreeMix` identifies gene flow into the outgroup; in this case, we place the root directly above this admixture node, as shown in Figure S2a, S5a, S6a.

##### 1.8 Model M2

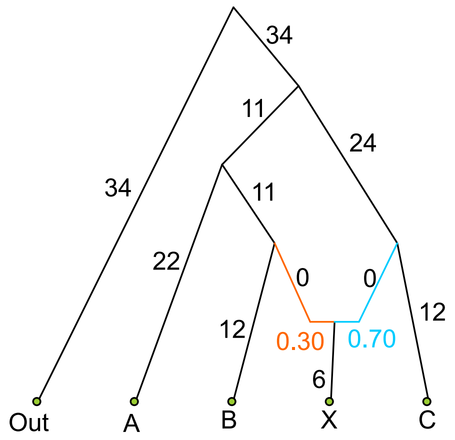

Figure S1: M2 based on Figure 5 of 4

#### 2 Supplementary Results

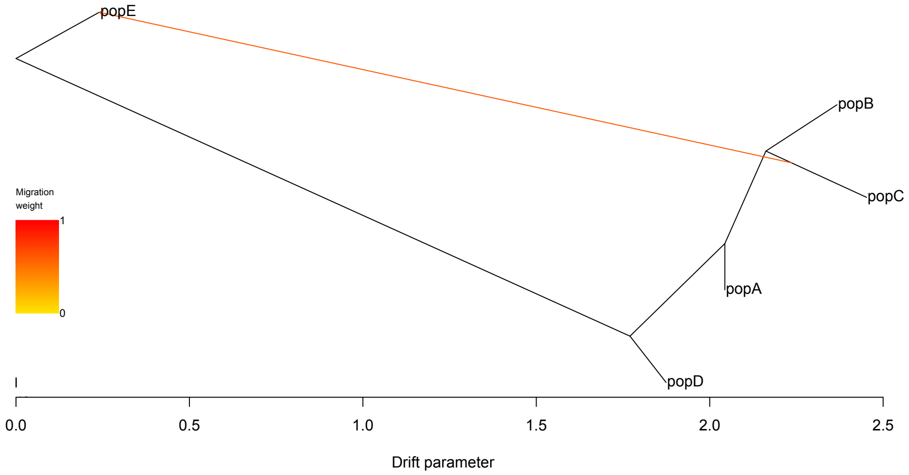

(a) `TreeMix` (log-lik = -366944; triplet dist = 9)

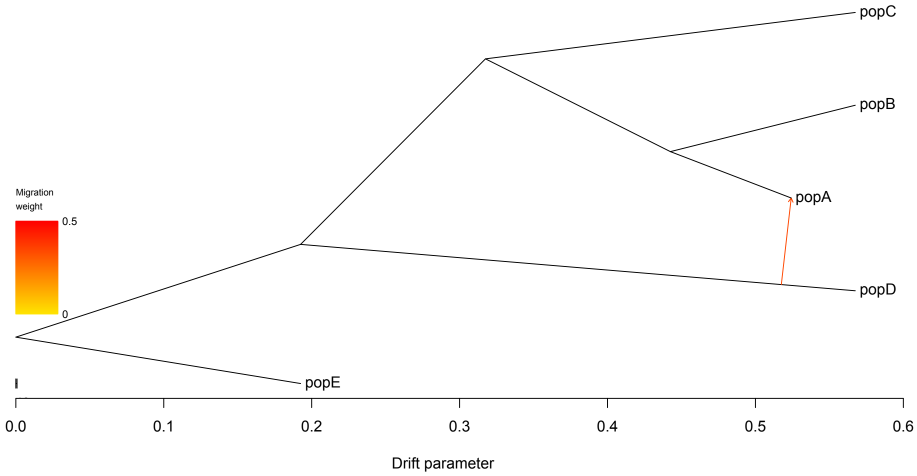

(b) `OrientAGraph (MLNO only)` (log-lik = 83; triplet dist = 0)

Figure S2: On M1, **`TreeMix`** finds an incorrect directed admixture graph topology, but **`OrientAGraph`** (with **`-mlno`** option only) finds the true one. There is a difference in the log-likelihood scores of these two graphs.

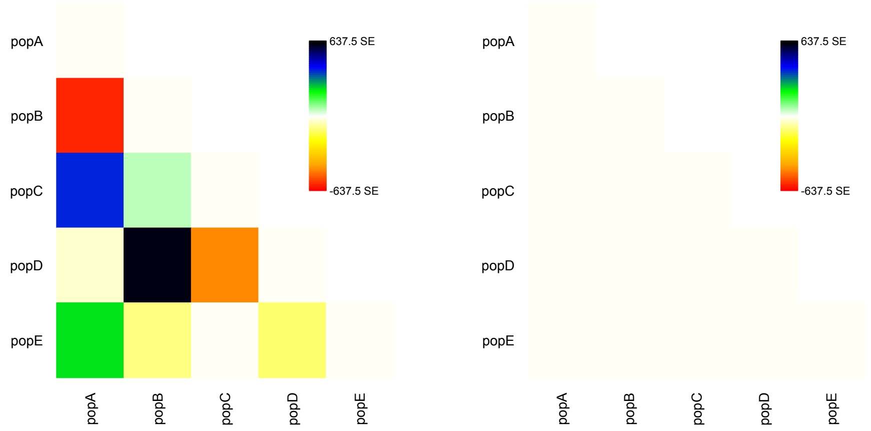

(a) `TreeMix` (b) `OrientAGraph (MLNO only)`

Figure S3: On M1, `TreeMix` finds an incorrect directed admixture graph topology, but `OrientAGraph` (with `-mlno` option only) finds the true one. There is a difference in residuals of these two estimated admixture graphs. Note that the standard error of all $f_2$-statistics was set to 0.0001.

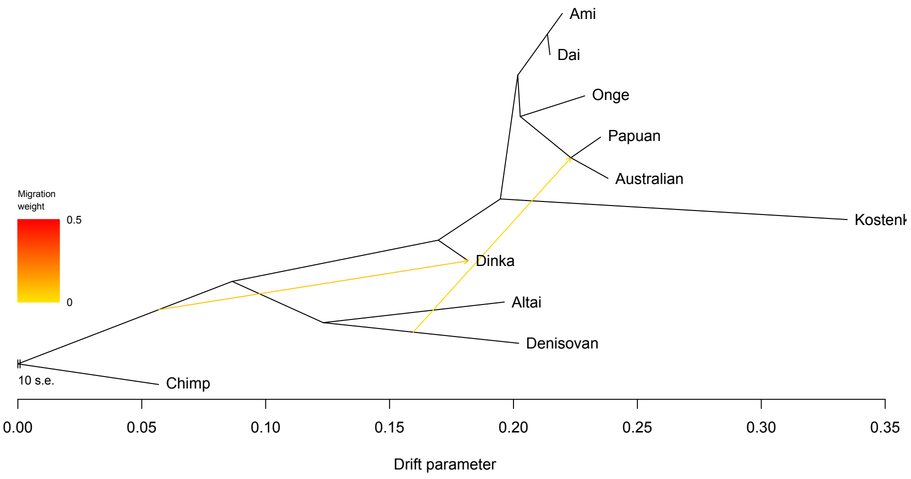

(a) TreeMix (log-lik = 137; triplet dist = 14)

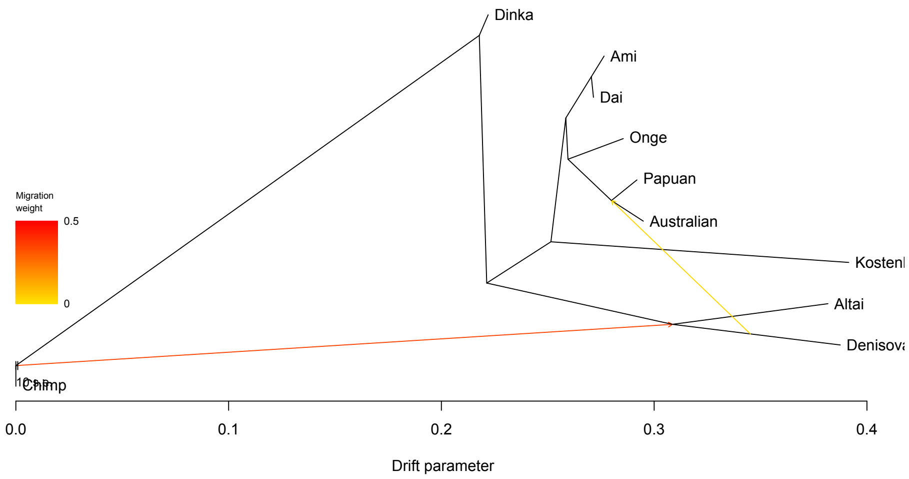

(b) OrientAGraph (MLNO only) (log-lik = 137; triplet dist = 14)

Figure S4: On M4, **TreeMix** and **OrientAGraph** (with **-mlno** only) recover different directed graph topologies for different population orders. All of the topologies returned were incorrect and had a triplet distance of 14 to the true admixture graph. **TreeMix** and **OrientAGraph** (with **-mlno** only) returned a graph with a log-likehood score of 137 on 60% and 86% of population orders, respectively. Some graphs returned by these methods with these scores are shown above.

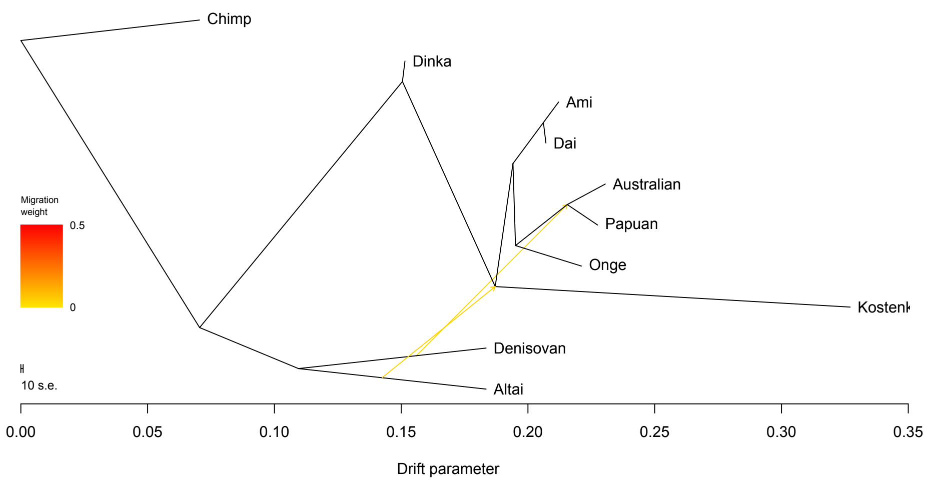

`OrientAGraph (ALLMIGS only)` (log-lik = 373; triplet dist = 0)

Figure S5: On M4, **`OrientAGraph`** (with **`-allmigs`** only or with **`-allmigs -mlno`** options) recovers true directed admixture graph topology. This graph has a log-likelihood score of 373, whereas the graphs found by **`TreeMix`** or **`OrientAGraph`** (with **`-mlno`** option) had log-likelihood scores of 137.

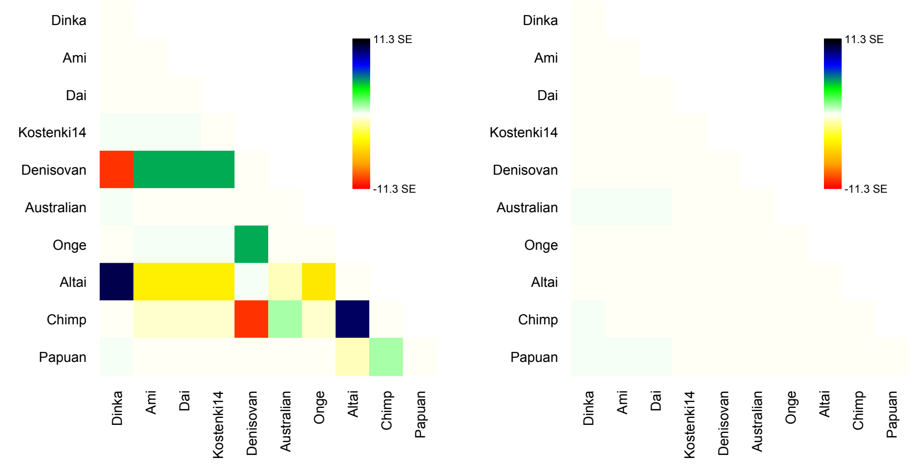

(a) `TreeMix` (b) `OrientAGraph (ALLMIGS only)`

Figure S6: On M4, **`TreeMix`** finds an incorrect directed admixture graph topologies, but **`OrientAGraph`** (with **`-allmigs`** only or with **`-allmigs -mlno`**) finds the true one. There is a difference in residuals of the estimated graphs. The residuals shown here are for the trees plotted in Figure S4a (**`TreeMix`**) and Figure S5 (**`OrientAGraph`**with **`-allmigs`**), respectively. Note that the standard error of all $f_2$-statistics was set to 0.0001.

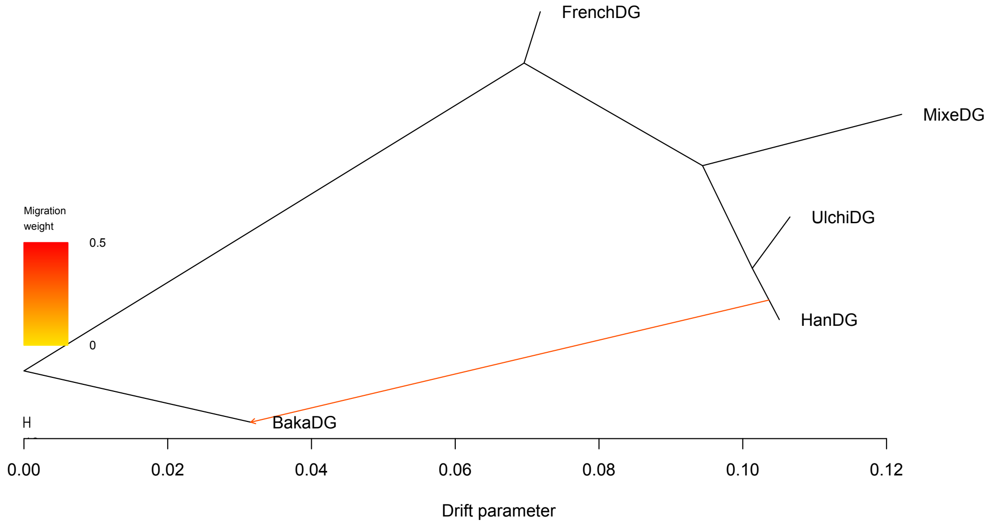

(a) `TreeMix` (log-lik = -33; triplet dist = 9)

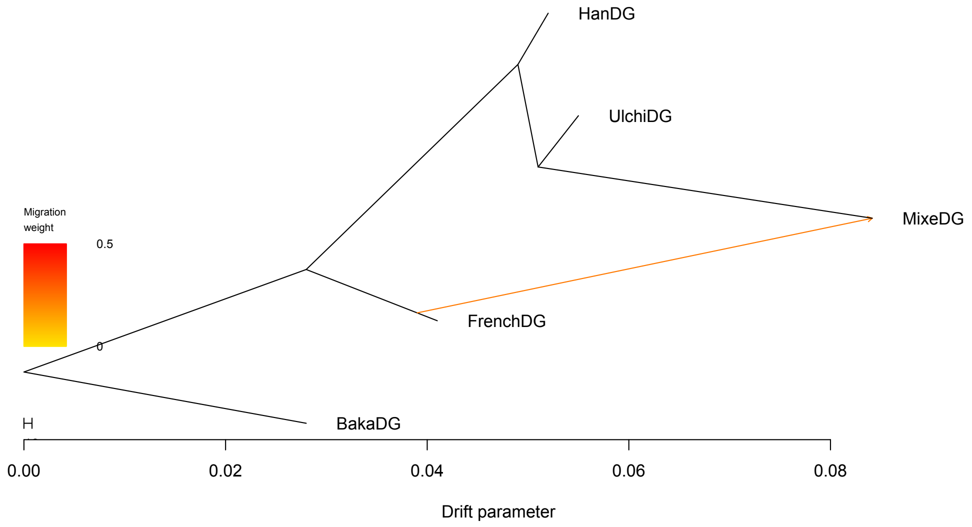

(b) `OrientAGraph (MLNO only)` (log-lik = 83; triplet dist = 0)

Figure S7: On M5, TreeMix finds an incorrect directed admixture graph topology, but `OrientAGraph` (with `-mlno` option only) finds the correct one. There is a difference in the log-likelihood scores of these two graphs.

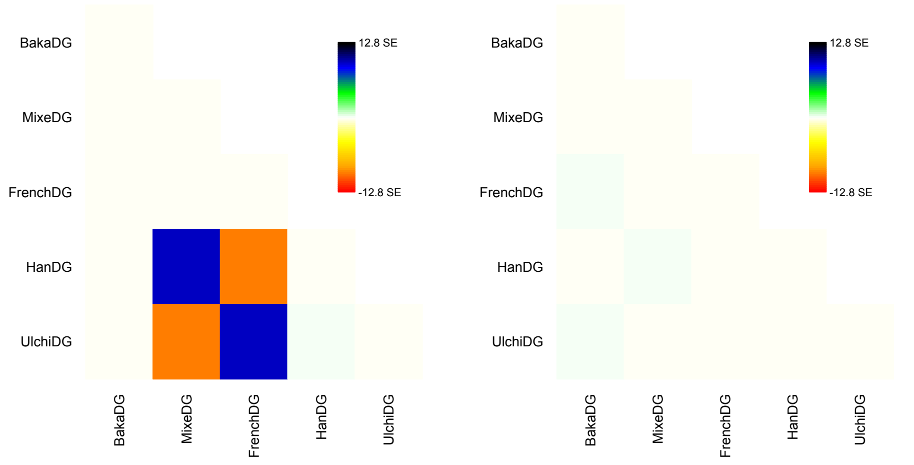

(a) `TreeMix` (b) `OrientAGraph (MLNO only)`

Figure S8: On M5, **`TreeMix`** finds an incorrect directed admixture graph topologies, but **`OrientAGraph`** (with **`-mlno`** option only) finds the true one. There is a difference in residuals of the estimated graphs. Note that the standard error of all $f_2$-statistics was set to 0.0001.

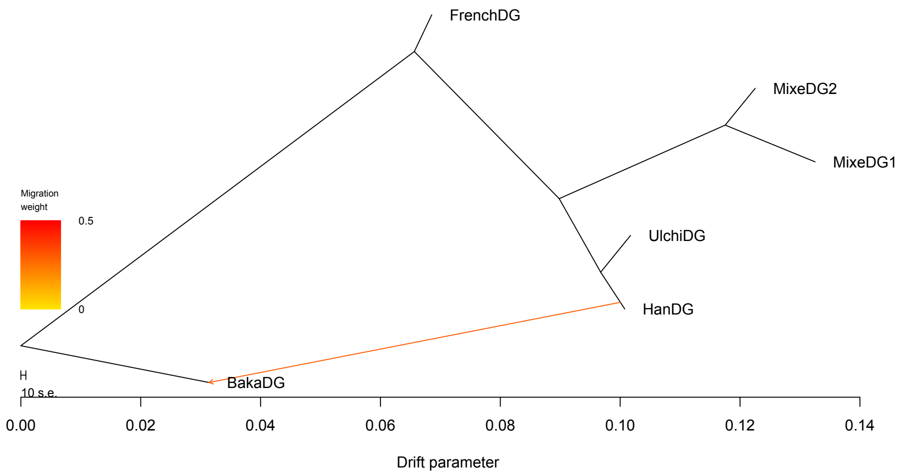

(a) `TreeMix` (log-lik = -30; triplet dist = 15)

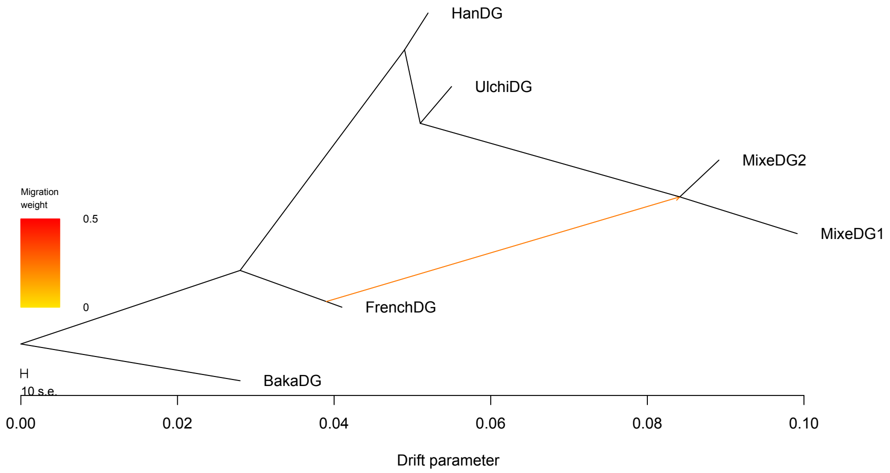

(b) `OrientAGraph (MLNO only)` (log-lik = 124; triplet dist = 0)

Figure S9: On M6, **`TreeMix`** finds an incorrect directed admixture graph topology, but **`OrientAGraph`** (with **`-mlno`** option only) finds the true one. There is a difference in the log-likelihood scores of these two graphs.

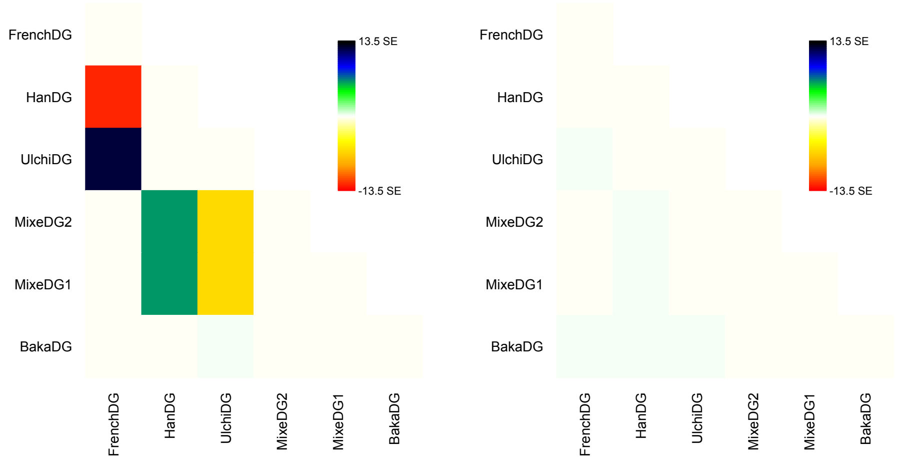

(a) `TreeMix` (b) `OrientAGraph (MLNO only)`

Figure S10: On M6, `TreeMix` finds an incorrect directed admixture graph topology, but `OrientAGraph` (with `-mlno` option only) finds the true one. There is a difference in residuals of these two estimated admixture graphs. Note that the standard error of all $f_2$-statistics was set to 0.0001.

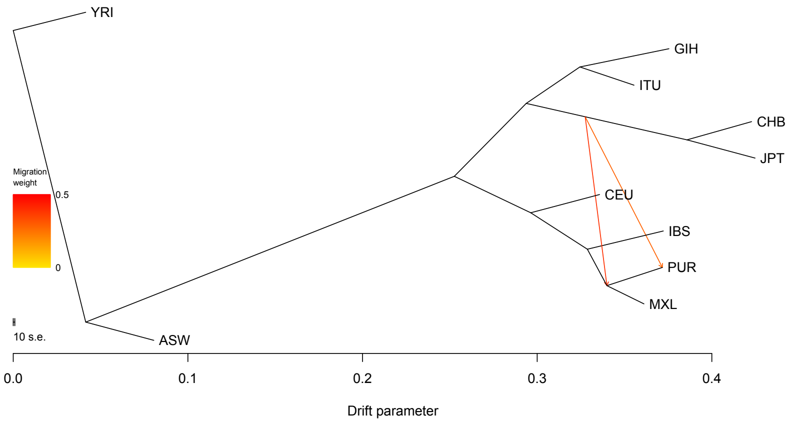

(a) `TreeMix` (log-lik = -2883; triplet dist = 20)

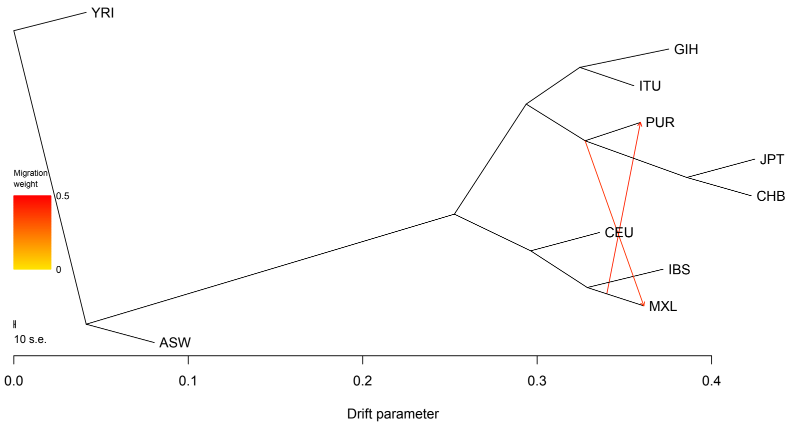

(b) `OrientAGraph (MLNO only)` (log-lik = -2883; triplet dist = 16)

Figure S11: On M7, we ran `TreeMix` and `OrientAGraph`, both with 100 different population addition orders, selected uniformly at random. `TreeMix` returned a graph with triplet distance of 20 to the correct admixture graph on 67% of runs. `OrientAGraph` (MLNO only) returned a graph with triplet distance of 16 on 55% of runs. `OrientAGraph` (ALLMIGS only and MLNO+ALLMIGS) returned a graph with a triplet distance of 20 on 100% of runs. Both of these graphs seem to have the same base tree, but there is an edge of length 0 (it looks like there is a vertex with outdegree 3, which is not allowed by `TreeMix`'s data structure). Different resolutions in the branching order around this edge would explain why these graphs have the same likelihood score but slightly different triplet scores.

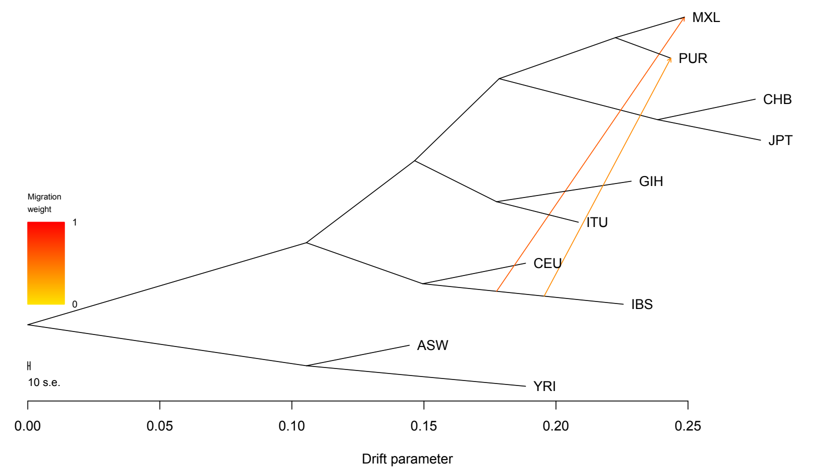

`True Graph scored by OrientAGraph` (log-lik = 373)

Figure S12: On M7, we used **`OrientAGraph`** to score the true admixture graph topology; the result is shown above.

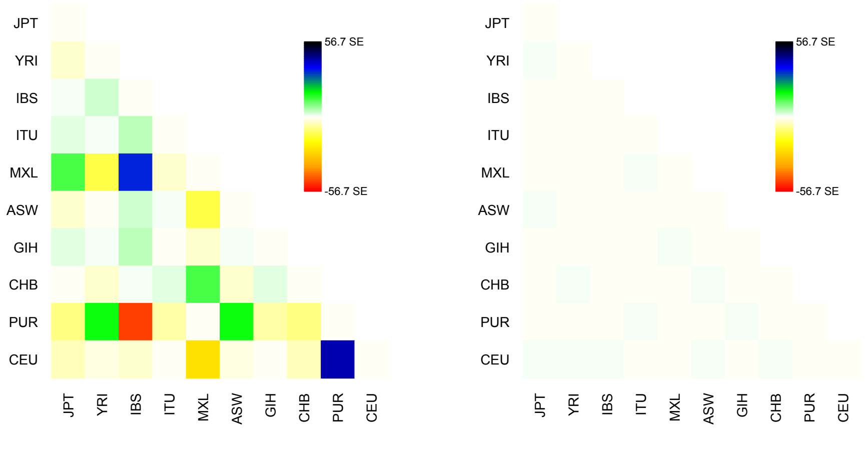

(a) `TreeMix / OrientAGraph`

(b) `True Graph (scored by OrientAGraph)`

Figure S13: On M7, **`TreeMix`** and **`OrientAGraph`** find incorrect directed admixture graph topologies with likelihood score of -2883. We gave **`OrientAGraph`** the true graph topology as input, so that it could fit its parameters and compute its likelihood score (373). There is a difference in residuals of these graphs. Note that the standard error of all $f_2$-statistics was set to 0.0001.

Table S1: Results for estimating admixture graphs on given the $f$-statistics implied by the models in Figure 3 in the main text. Admixture graphs were estimated using `TreeMix` as well as two versions of `admixturegraph` , one using the MLNO subroutine only and the other using an exhaustive search for the best admixture edge addition (ALLMIGS). We report the log-likelihood score, the topological error (i.e. the triplet distance) between the model admixture graph and the estimated admixture graph, and the running time in seconds. All methods were run 100 times, as discussed in Section 5 in the main text, so the running time is the average of 100 runs. *Indicates there was a difference in the log-likelihood score or triplet distance for the graphs returned on different runs, in which case, we report the average.

| Model | Log-likelihood | | | Topological Error | | | Running Time | | |
|---|---|---|---|---|---|---|---|---|---|
| | `TreeMix` | `OrientAGraph` | | `TreeMix` | `OrientAGraph` | | `TreeMix` | `OrientAGraph` | |
| | | ALLMIGS only | MLNO only | | ALLMIGS only | MLNO only | | ALLMIGS only | MLNO only |
| M1 | -366944 | -366944 | 83 | 9 | 9 | 0 | 0.06 | 0.13 | 0.11 |
| M2 | 83 | 83 | 83 | 0 | 0 | 0 | 0.04 | 0.11 | 0.08 |
| M3 | 124 | 124 | 124 | 0 | 0 | 0 | 0.28 | 0.64 | 1.83 |
| M4⋆ | 103* | 373 | 130* | 14 | 0 | 14 | 1.28 | 5.10 | 7.36 |
| M5 | -33 | -33 | 83 | 9 | 9 | 0 | 0.05 | 0.14 | 0.09 |
| M6⋆ | -30 | -30 | 124 | 15 | 15 | 0 | 0.11 | 0.25 | 0.17 |
| M7 | -2883 | -2883 | -2883 | 18.6* | 20 | 17.8* | 5.58 | 8.00 | 10.77 |
| M8 | 124 | 124 | 124 | 0 | 0 | 0 | 0.09 | 0.27 | 0.16 |

### References

[1] A. R. Francis and M. Steel. Which Phylogenetic Networks are Merely Trees with Additional Arcs? *Systematic Biology*, 64(5):768–777, 06 2015.

[2] K. T. Huber, L. van Iersel, R. Janssen, M. Jones, V. Moulton, Y. Murakami, and C. Semple. Rooting for phylogenetic networks. *arXiv*, CoRR:abs/1906.07430, 2019.

[3] J. Jansson, K. Mampentzidis, R. Rajaby, and W.-K. Sung. Computing the Rooted Triplet Distance Between Phylogenetic Networks. In C. J. Colbourn, R. Grossi, and N. Pisanti, editors, *Combinatorial Algorithms*, pages 290–303, Cham, 2019. Springer International Publishing.

[4] N. Patterson, P. Moorjani, Y. Luo, S. Mallick, N. Rohland, Y. Zhan, T. Genschoreck, T. Webster, and D. Reich. Ancient Admixture in Human History. *Genetics*, 192(3):1065–1093, 2012.

[5] J. K. Pickrell and J. K. Pritchard. Inference of Population Splits and Mixtures from Genome-Wide Allele Frequency Data. *PLOS Genetics*, 8(11):1–17, 11 2012.
